# IFT20 mediates trafficking of ciliary proteins to support mammalian motile ciliogenesis in testicular efferent ductules and respiratory tract

**DOI:** 10.64898/2026.11.12.729724

**Authors:** Tao Li, Alain Schmitt, Jiehong Pan, Cheng Zheng, Wei Li, Shahnaz Parveen, Anuradha Yadav, Mohammad Abdulghani, Kristian Bennett, Rex A Hess, Gregory J Pazour, Christopher V Kelly, Donghui Huang, Steven L Brody, Aminata Touré, Zhibing Zhang

**Affiliations:** Institute of Reproduction Health Research, Tongji Medical College, Huazhong University of Science and Technology, Wuhan 430030, China; Department of Physiology, Wayne State University, Detroit, MI, 48201, USA; Université Paris Cité, INSERM U1016, CNRS UMR8104, Institut Cochin, Paris, France; Division of Pulmonary and Critical Care Medicine, Department of Medicine, Washington University School of Medicine, St. Louis, MO, 63110, USA; Department of Occupational and Environmental Health, School of Public Health, Wuhan University of Science and Technology, Wuhan, Hubei 430065, China; Department of Physics and Astronomy, Wayne State University, Detroit, MI, USA; Department of Comparative Biosciences, College of Veterinary Medicine, University of Illinois, 2001S. Lincoln, Urbana, IL, USA; Program in Molecular Medicine, University of Massachusetts Chan Medical School, Worcester, MA, USA; Université Grenoble Alpes, INSERM U1209, CNRS UMR 5309, Team Physiology and Pathophysiology of Sperm cells, Institute for Advanced Biosciences, 38000 Grenoble, France; Department of Obstetrics & Gynecology, Wayne State University, Detroit, MI, 48201, USA

## Abstract

Cilia are antenna-like organelles that extend from the apical cell membrane and are broadly present in eukaryotes. They are classified as primary and motile types; the later including sperm flagellum and somatic multiciliated cells present within the reproductive and respiratory systems. Intraflagellar transport (IFT) provides delivery of cargo for cilia assembly and maintenance. Of the multiple protein components, IFT20, a component of the intraflagellar transport-B (IFT-B) complex, is required for cargo delivery anterograde during primary cilia and mouse sperm flagellum formation, and is also known to link the Golgi with the primary cilia; however, its role in motile cilia from somatic cells is so far unknown. The objective of this study is to investigate, in mice, the role of IFT20 in motile cilia of somatic cells. We used a FOXJ1-Cre mouse to conditionally delete Ift20 only in motile cilia. We found that most *Ift20* conditional knock out (cKO) mice were viable and none showed left-right asymmetry defects nor hydrocephalus, though they exhibited reduced body weight. Compared to control mice, adult cKO males were subfertile with significantly shortened motile cilia and decreased ciliary beating frequency (CBF) in efferent ductules (EDs). Examination of ciliary ultrastructure in EDs revealed partial absence or displacement of the peripheral doublet microtubules (DMTs), as well as defects in both the inner and outer dynein arms and the central pair apparatus. In addition, the lumen of EDs were significantly dilated and filled with aggregated sperm. Sperm aggregation also led to a dilatation of the lumen of the seminiferous tubules and testicular atrophy, accompanied by an increased number of degenerated germ cells that were sloughed into the lumen. The tracheal cilia were minimally shortened and had reduced CBF, however, they had normal appearing ultrastructure. In contrast to control mice, analysis of cKO ciliated cells, in both EDs and trachea, showed an accumulation of the central apparatus protein SPAG16, SPAG17, the outer dynein arm DNAI2, and inner dynein arm DNAH2 in the apical region of the cytoplasm. These data indicate that IFT20 is essential in sperm and somatic ciliated cells from ED and trachea, to sustain flagella/cilia formation and beat frequency and suggest that, IFT20 assists in trafficking ciliary structures from the cytoplasm to the motile axoneme.

## Introduction

Cilia are hair-like organelles that project from the cell surface and have a microtubule-based structure. Based on their structure and function, they are classified into two types: primary cilia and motile cilia [**1**]. Primary cilia typically exist as solitary structures on most cells, characterized by a classic 9+0 microtubule arrangement, and are generally non-motile. They function in regulating cell growth, development, and metabolism by sensing the extracellular environment and mediating signal transduction [**2**]. In contrast, most motile cilia exhibit a typical 9+2 microtubule arrangement, consisting of a central pair of microtubules surrounded by nine peripheral microtubule doublets. They propel cell movement through rhythmic beating (e.g., sperm flagella) or regulate fluid flow over cell surfaces, as observed in the embryonic node, brain ependyma, airways, fallopian tube, and EDs of the testes [**3**]. Proteins and associated protein complexes required for ciliary function are synthesized in the cytoplasm, and then transported into the cilia via intraflagellar transport (IFT) machinery [**4**]. This mechanism is mainly mediated by two multi-protein complexes, IFT-A and IFT-B, each linked to cytoplasmic motor proteins (dynein-2 and kinesin-2, respectively [**5,6**]) to form cargo-carrying “trains”. IFT-B transports protein anterograde (from the cilia base to the tip) while IFT-A transports proteins retrograde (from the tip to the base).

The structure of IFT-A complex contains core (IFT144/IFT140/IFT122) and non-core (IFT139/IFT121/IFT43). The IFT-B complex consists of IFT20, IFT22, IFT25, IFT27, IFT38, IFT46, IFT52, IFT54, IFT56, IFT57, IFT70, IFT74, IFT80, IFT81, IFT88, and IFT172, and also comprises two subcomplexes. The IFT-B1 subcomplex contains a core structure centered on IFT52 that binds IFT88, IFT70, and IFT46and recruits the subcomplex IFT81-IFT74-IFT27-IFT25-IFT22 [**4**]. While IFT56 has been included in the IFT-B1 subcomplex due to its interaction with IFT46 [**7**], it is established that mutations or deficiencies in IFT56 lead to shortened and/or structurally disorganized cilia [**8,9**]. The IFT-B2 subcomplex is formed by three dimers: IFT172–IFT80, IFT57– IFT38, and IFT54–IFT20 [**6,10–12**]. IFT trafficking is cyclical, characterized by distinct anterograde and retrograde phases [**4**]. Transport initiates at the basal body, where the IFT-A/B protein complexes first assemble into anterograde train structures. These trains load cargo proteins required for ciliary assembly and are subsequently translocated through the transition zone to the ciliary tip by kinesin motors. IFT train move to the tip, then, dissociate from the kinesin motors, and are remodeled into retrograde train structures [**13**]. These retrograde trains bind new cargo, associate with dynein-2 motors, and are then transported back toward the cell body [**4,5**].

IFT20, a component of the IFT-B protein complex, is the smallest protein (15 kDa) and, as the smallest IFT subunit, participates in the bidirectional trafficking of the IFT-A/B complex [**14**]. In contrast to other IFT proteins, IFT20 exhibits a dual subcellular localization. Beyond its presence at the ciliary basal body, it is also anchored to the Golgi apparatus via the protein GMAP210 [**15,16**] and facilitates the transport of proteins essential for primary cilium formation from the Golgi apparatus to the basal body of the primary cilium. The GM130-VPS15 protein complex regulates the release of IFT20-associated vesicles, which are rich in cilia membrane proteins [**17**], from the Golgi apparatus to the primary and motile cilia via the microtubule-based motor protein KIFC1 [**18–20**]. Subsequently, these IFT20-associated vesicles fuse with vesicles containing Rab8 and its effector Rabaptin5, a process facilitated by IFT54 (known as elipsa in zebrafish) [**21**]. Finally, Rab8 mediates the fusion of this composite vesicle with the plasma membrane at the ciliary base [**22**]. Furthermore, IFT20 participates in the initial stages of primary cilium formation. During centriole maturation and its transition into the basal body, the proteins EHD1/EHD3 recruit transition zone proteins along with IFT20. Depletion of EHD1/EHD3 leads to the accumulation of IFT20 at the base of the basal body and results in ciliogenesis defects [**23**]. Early ciliogenesis protein CCDC41 localizes to the distal end of the mother centriole and is anchored to the Golgi apparatus in an IFT20-dependent manner. Depletion of CCDC41 impairs the recruitment of IFT20 to the centrosome, while depletion of either CCDC41 or IFT20 inhibits ciliogenesis at the ciliary vesicle docking step [**24,25**]. Hence IFT20 not only functions as a subunit of the IFT-B complex for the IFT process but is involved in the initial phase of ciliogenesis and subsequent membrane protein transport from the cytoplasm to cilia.

Our prior work has especially focus on the sperm flagellum, a specialized type of motile cilium, that forms during the late stages of germ cell development [**26**]. One line of investigation into IFT20 function in motile cilia stems from our previous study using germ cell-specific conditional knockout (cKO) mouse model [**27**]. Sperm from IFT20 cKO mice exhibited reduced motility, accompanied by shortened and coiled flagella. Transmission electron microscopy revealed the loss of the typical “9+2” microtubule arrangement in the flagella, indicating that IFT20 is essential for sperm flagellum formation, likely via the IFT machinery [**27**]. However, it remains unclear whether the function of IFT20 is conserved in other motile cilia.

In particular, we sought to focus on an understudied specialized structure rich in motile cilia that is part of the male reproductive system—the Efferent Ductules (EDs). These structures connect the rete testis to the epididymis. Following their release from the seminiferous tubules, spermatozoa are collected within the rete testis and subsequently transported through the EDs to the caput epididymis [**3,28**]. Within EDs, cilia function as agitators, generating vigorous fluid turbulence to prevent sperm aggregation during transport [**29**]. and studies of mouse mutant models have shown that inadequate ED ciliary motility increases pressure within the seminiferous tubulesleading to progressive testicular atrophy over time [**29**]. Our previous study [**30**] was the first to demonstrate that loss of the IFT-A complex protein IFT140 disrupts IFT trafficking in the ED, as well as the airway epithelia, leading to cytoplasmic cargo accumulation and consequently impaired motile ciliogenesis and male infertility. However, the functional impact of IFT-B subunits on mammalian motile cilia remains unclear. To investigate its functional role in mammalian motile cilia, we employed *FOXJ1*-Cre to specifically delete the IFT-B subunit IFT20 in motile cilia-bearing cells in mice.

## Materials and Methods

### Ethics statement

All animal research was executed in compliance with the guidelines of the Wayne State University Institutional Animal Care and the Program Advisory Committee (Protocol number: 24-02-6561).

### Generation of *Ift20^flox/flox^; FOXJ1*-Cre mouse model

Floxed *Ift20* mice were provided by Dr. Gregory J. Pazour, University of Massachusetts Medical School [**31**]. The *FOXJ1*-Cre transgenic mouse lines were sourced from Dr. Michael J. Holtzman at the Washington University School of Medicine in St. Louis [**32**]. Floxed *Ift20* homozygous mice were bred with *FOXJ1*-Cre mice to obtain first-generation (F1) mice heterozygous for *Ift20* and positive for *FOXJ1*-Cre. These F1 mice were then crossed with homozygous floxed *Ift20* mice to generate floxed *Ift20* homozygous mice that were also *FOXJ1*-Cre-positive conditional knockout (cKO) mice.

### RT-PCR

Total RNA was extracted from various organs of adult mice using TRIzol reagent (Thermo Fischer Scientific). Subsequently, reverse transcription was performed with a reverse transcription kit (Maxima H Minus First Strand cDNA Synthesis Kit, K1651). The obtained cDNA was diluted 10-fold and subjected to conventional PCR amplification. All PCR products were then separated by agarose gel electrophoresis, and experimental results were recorded following imaging.

### Fertility Assessment

To evaluate male fertility, adult cKO mice (2-4 months old) and control littermates were co-housed with healthy, reproductively competent wild type female mice (2-4 months old) in a 1:1 pairing ratio. Breeding pairs were continuously monitored for 2 months, with recording of the total number of litters per female and pup count per litter.

### Tissue histology

Testes, efferent ducts, epididymides, and trachea were dissected and fixed overnight at 4°C in 4% Formalin fixative. The tissues were subsequently embedded in paraffin, and 5 μm sections were prepared using a microtome, followed by standard hematoxylin and eosin (H&E) staining procedures. Slides were examined using a BX51 Olympus microscope (Olympus Corp., Melville, NY, Center Valley, PA), and photographs were taken with a ProgRes C14 camera (Jenoptik Laser, Germany).

### Cilia beat frequency measurement

#### Sample preparation

For testicular EDs, immediately after isolation from the mouse, the tissue was immersed in pre-warmed (37°C) HEPES-Ringer solution. Following adipose tissue removal, the ductules were longitudinally incised and transferred to a glass slide. A sufficient volume of HEPES-Ringer solution was applied to maintain hydration. Two side supporting coverslips (1-2 per side) were positioned adjacent to the tissue to create a spacer, and a final coverslip was gently placed over the sample. This configuration ensured mechanical stability and adequate space for ciliary motility. The prepared specimen was then mounted on a thermostatically controlled microscope stage (37°C), and videos were captured using a 20× objective. For tracheal samples, the trachea was transversely sectioned to prepare ring-shaped tracheal segments, facilitating optimal lateral exposure of motile cilia. During slide preparation, additional coverslips (2-3 per side) were positioned as spacers, while other procedures remained identical to those described for testicular EDs. The imaging system comprises an Olympus inverted microscope (OLYMPUS CKX41) and a Basler high-speed camera (Basler acA1300-200μm).

#### CBF measurement

CBF was quantified as described by Horani et al[**33**], using the Sisson-Ammons Video Analysis (SAVA, Ammons Engineering, v2.1.2w). All motile cilia observed under the microscope were included in the CBF analysis. CBF measurements were performed as follows: for each ED, ≥10 video recordings were acquired, with 8-10 regions of interest (ROI) analyzed per video, and for trachea sample, ≥5 videos were recorded, and 3–5 ROI per video were analyzed for data acquisition.

### Immunofluorescence staining

Mouse trachea and EDs were fixed in 4% paraformaldehyde (PFA) were embedded in paraffin, and 5 μm sections were prepared. After dewaxing and rehydration, antigen retrieval was performed by heating the sections in sodium citrate buffer (pH 6.0) using a microwave oven for 20 min. The sections were then washed twice with 1× PBS (5 min per wash). Non-specific binding sites were blocked with 10% goat serum in a humidified chamber at room temperature for 1 hour. Primary antibodies diluted in 10% goat serum were applied to the sections and incubated overnight at 4°C in a humidified chamber. The following day, sections were removed from 4°C, and the primary antibody solution was gently removed. Sections were washed three times with 1× PBS (10 min per wash at room temperature). Excess liquid was carefully blotted, and species-matched secondary antibodies (appropriately diluted) were applied to the sections. Incubation proceeded for 1 hour at room temperature in a light-protected humidified chamber. After secondary antibody removal, sections were washed three times with 1× PBS (10 min per wash at room temperature) under ambient light protection. Sections were then briefly dried, and antifade mounting medium containing DAPI was applied to the tissue areas. Coverslips were mounted, and nail polish was used to seal the edges to prevent desiccation. Fluorescence images were acquired using a Carl Zeiss Axio Imager M2 OEM 01 fluorescence microscope equipped with a Dhyana 400D camera, and subsequent image merging was performed with ImageJ software. The antibodies used and their details are provided in Supplementary Table 1.

### TUNEL Analysis

Paraffin-embedded, 5 μm thick sections were selected and processed for routine dewaxing and hydration. Subsequent steps were performed according to the instructions of the In Situ Cell Death Detection Kit (Roche, 11684795910). Briefly, antigen retrieval was carried out using sodium citrate buffer (pH 6.0) under microwave heating for 2 min, followed by two 5-min washes with PBS. During this period, the working solution was prepared by mixing the enzyme solution and the label solution at a 1:9 ratio. The resulting working solution was applied to the tissue sections and incubated in a dark, 37 °C humidified chamber for 1 h. After incubation, the sections were washed three times with PBS in the dark for 5 min each. Finally, the sections were mounted using an anti-fade mounting medium containing DAPI. The results were observed and recorded under a fluorescence microscope, as described for Immunofluorescence Staining.

### Electron microscopy

Mouse EDs and trachea were fixed by incubation in 0.1 M phosphate buffer (pH 7) supplemented with 3% glutaraldehyde (Grade I; Sigma-Aldrich) for 2 h at room temperature. The samples were washed twice in PBS and resuspended in 0.2M sodium cacodylate buffer. The samples were then post-fixed by incubation with 1% osmium tetroxide (Electron Microscopy Sciences), after which they were dehydrated by immersion in a graded series of alcohol solutions and embedded in Epon 812 resin (Electron Microscopy Sciences). Semi-thin sections were cut and stained with toluidine blue-Azur II. Ultra-thin sections (90 nm) were cut with a Leica UC 7 ultramicrotome (Leica Microsystems and were then stained with uranyl acetate and lead citrate. Sections were analyzed with a JEOL 1011 microscope and digital images were acquired with a Gatan Orius 1000 CCD camera and Digital Micrograph software.

### Cilia length measurement

Based on the results of HE stains, the length of motile cilia in the testicular efferent ducts and tracheal epithelium was measured using the measurement tool in the software ImageScope (v.12.4.6.5003). The data were analyzed statistically using GraphPad Prism (v9.0.0.121).

### Protein Extraction and Western Blot Analysis

Testicular EDs were aseptically dissected from mice and homogenized in RIPA lysis buffer supplemented with a protease inhibitor cocktail. Protein concentrations were quantified using a bicinchoninic acid (BCA) assay kit (BIO-RAD), followed by preparation of loading buffer containing equal protein amounts for each sample. Proteins were separated via sodium dodecyl sulfate-polyacrylamide gel electrophoresis (SDS-PAGE). Post-electrophoresis, gels were transferred onto polyvinylidene difluoride (PVDF) membranes under chilled conditions (4°C). Membranes were blocked for 1 hour at room temperature with gentle agitation in Tris-buffered saline containing 0.1% Tween-20 (TBST) and 5% non-fat dry milk. Primary antibodies (diluted in 5% milk-TBST) were applied to membranes and incubated overnight at 4°C with continuous shaking. Next, membranes were washed three times in TBST (10 min per wash, room temperature with agitation). Species-matched horseradish peroxidase (HRP)-conjugated secondary antibodies (diluted in 5% milk-TBST) were then applied and incubated for 1 hour at room temperature with shaking. After three additional TBST washes (10 min each), membranes were incubated with enhanced chemiluminescence (ECL) substrate (mixed 1:1 from components A and B) for 1 minute. Signal detection was performed using a BIO-RAD ChemiDoc MP Imaging System. The antibodies used and their details are provided in Supplementary Table 1.

### Statistical analysis

Graphs were created using Microsoft Excel and GraphPad Prism (v9.0.0.121). Statistical analyses were performed using Student’s t-test. P > 0.05 was considered as not significant. By convention, *p < 0.05, **P<0.01, ***P<0.001, ****P<0.0001.

## Result

### IFT20 is highly expressed in the EDs, and its absence results in male infertility

It is well established that mouse *Ift20* plays a key role in primary cilia formation, and we previously reported that mouse IFT20 is essential for sperm formation. To determine if IFT20 is important for motile ciliogenesis in other somatic tissues, we first profiled protein levels across different organs. IFT20 was highly expressed in the testis, kidney, ovary, and EDs, while, lower levels of expression are found in the heart, liver, and brain (**Figure 1A**). To investigate the role of IFT20 in motile cilia, we crossed *Ift20*^flox/flox^ homozygous mice[**31**] with *FOXJ*-Cre mice, to specifically delete *Ift20* in motile cilia-bearing cells. In this study, mice with the genotype *FOXJ*-Cre^+^; *Ift20*^flox/flox^ are referred to as conditional knockout (cKO) mice (**Supplementary Figure 1A**). Genotyping was performed by PCR (**Supplementary Figure 1B**). Knockout efficiency in the EDs was confirmed by western blot (**Figure 1B**) and immunofluorescence staining (**Figure 1C**); in both experiments, the IFT20 protein was absent in the EDs of cKO mice. At one week of age, no significant difference in body weight was observed between cKO and control mice (**Figure 1D**). However, from the second week onward, cKO mice exhibited lower body weight compared to controls, a trend that persisted into adulthood (**Figure 1D, E, Supplementary Figure 2**). During our study, no control mice died (n>100); however, two homozygous cKO mice (n=42) died for unknown reasons. Fertility tests indicated a significant decline in the fertility of cKO male mice, with most being completely infertile (**Figure 1F, G**). As motile cilia are also present in the oviducts. We tested the fertility of the homozygous females, among the six breeding pairs, only one exhibited infertility.

**Figure 1.**
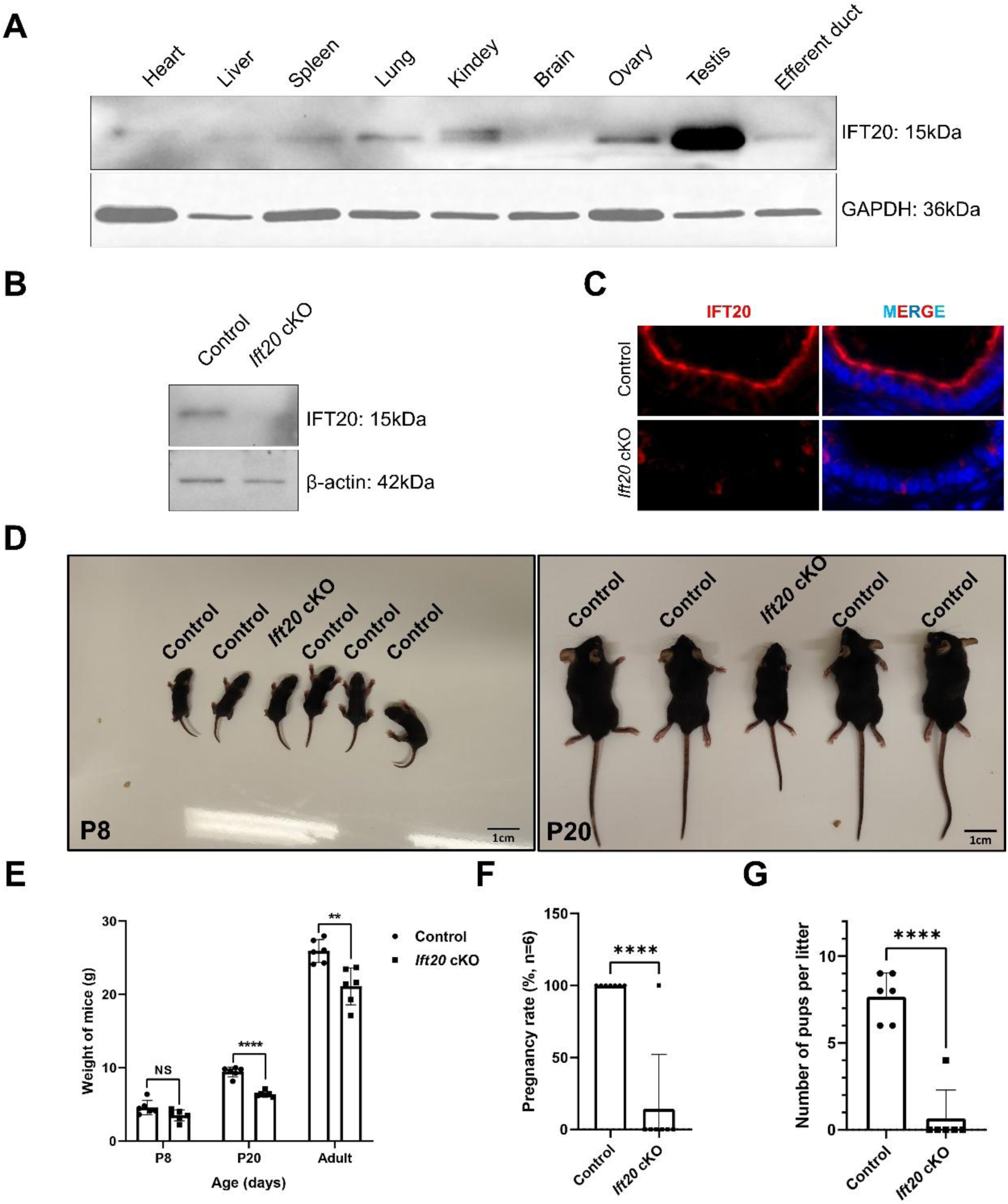
IFT20 is essential for the ciliary beating of the EDs and male fertility. **A**. Examination of IFT20 protein expression in mouse tissues by Western blot analysis. IFT20 was detected in most tissues, including the efferent duct. No signal was detected in the brain. Examination of knockout efficiency in the EDs of *Ift20* cKO mice. Western blot (**B**) and immunofluorescence staining (**C**) were conducted. IFT20 is missing in the cKO mice. **D**. *Ift20* cKO mice showed growth retardation. A representation image showing the same size of the P8, P20 control and cKO mice. **E**. Statistical analysis of the body weight of 8-day-old, 14-day-old, 20-day-old, and adult control mice and cKO mice. There was no significant difference at 8-day-old. After two-weeks-old, the cKO mice showed significantly lower body weight. **F**. Reduced fertility of *Ift20* cKO males Fertility rate of control and cKO mice. The fertility rate was significantly lower in the *Ift20* cKO mice. **G**. Litter size of control and cKO mice. The litter size was significantly lower in the *Ift20* cKO mice. n=6. *P<0.05; ** P<0.01; **** P<0.0001 compared to the control group.

Motile cilia are also present on the ependymal lining of the brain ventricles. Similar to those in the airways, they exhibit coordinated, directional beating to regulate cerebrospinal fluid (CSF) circulation [**34**]. However, in cKO mice, gross anatomical examination revealed no obvious macroscopic differences in brain morphology, and ventricular size was similar to controls **(Supplementary Figures 3, 4)**. This finding aligns with the results we previously reported for the *Ift140* knockout model [**30**]. No situs inversus was found in all cKO mice analyzed in the study (n=42). Histology of major organs, including the kidney and liver, was examined in the control and *Ift20* cKO mice. In mice with motile cilia defects, no cystic changes were found in these organs **(Supplementary Figure 4).**

### Disruption of *Ift20* in the EDs affected motile cilia formation and function

To investigate the overall effect of disruption of *Ift20* in the male reproductive organs, the testis, EDs, and epididymis were analyzed. Compared to the control mice, most cKO mice exhibited dilated efferent ducts and a more transparent, flattened epididymis throughout the caput, corpus, and cauda, suggesting impaired sperm transport transit the efferent ducts (**Figure 2A**).

**Figure 2.**
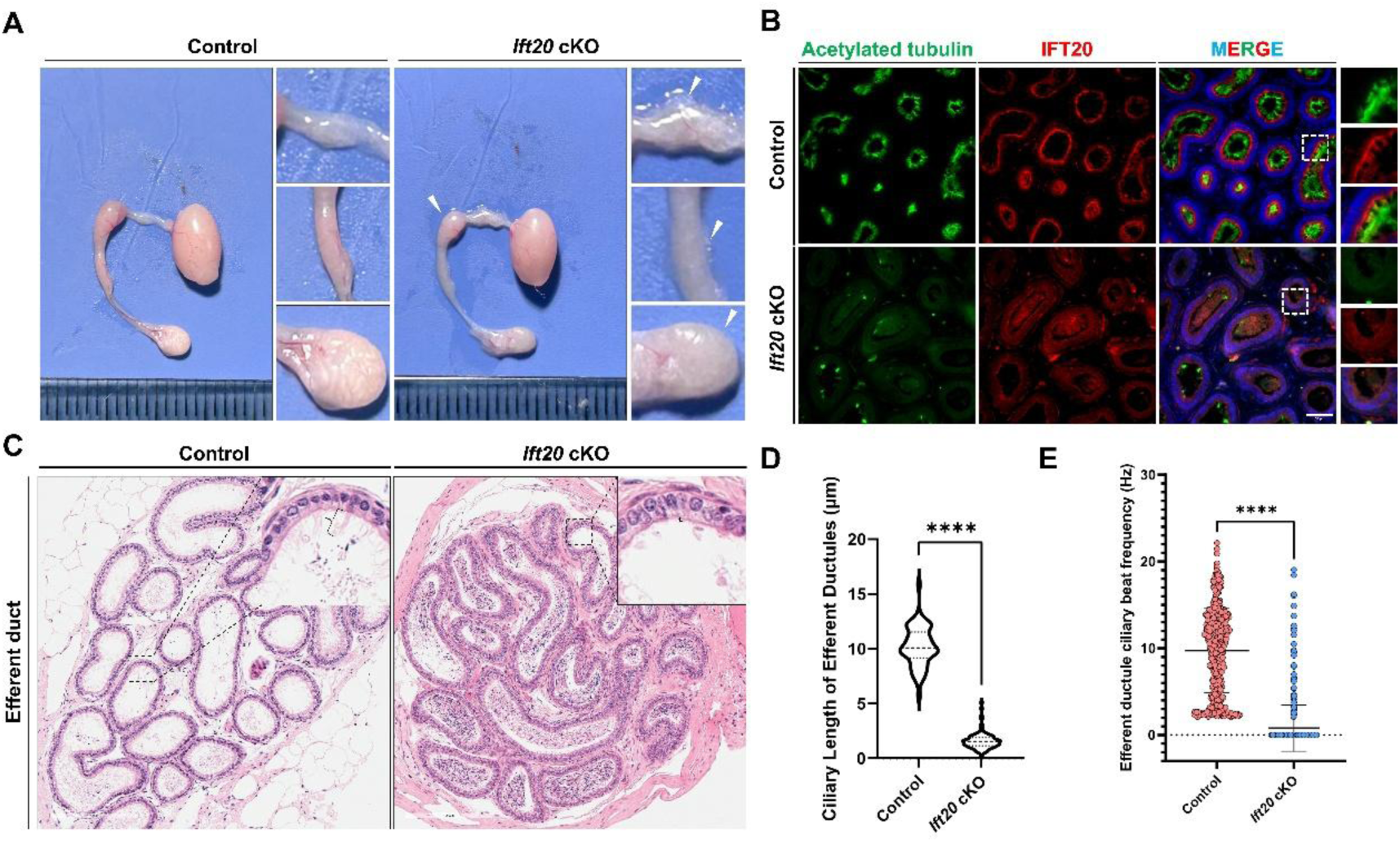
Shortened Length and Reduced Beating Frequency of Motile Cilia in the Efferent Ducts of cKO Mice **A**: Representative photomicrographs of the testis, efferent ducts, and epididymis from 4-month-old mice. Compared to tissues from control mice, the cKO mice exhibited dilated efferent ducts and a more transparent, flattened epididymis in carput, corpus, and cauda, suggesting impaired sperm transport through the efferent ducts (white arrows). **B**: Immunofluorescence was used to detect the knockout efficiency of IFT20 and the presence of motile cilia. Tissue from 4-month-old mice shows loss of IFT20 signal in the efferent ducts of cKO mice, confirming the absence of IFT20. Acetylated tubulin signal is absent in the vast majority of tubules, indicating loss of motile cilia. The scale is in the corner. **C**: Histological examination with H&E staining of the samples in panel A revealed that, compared to the control mice, the cKO mice exhibited the lumina of the efferent ducts showed evident sperm accumulation, accompanied by a marked shortening of the motile cilia. The scale is in the corner. **D**: The length of motile cilia in the efferent ducts was measured using samples from three pairs of mice. A total of 54 ducts were analyzed in the control group (mean 10.08 µm), while 110 ducts were included from the cKO mice (mean 1.52 µm). The results indicate a significant shortening of motile cilia in the cKO mice. **** P<0.0001 compared to the control group. **E**: The ciliary beat frequency in the efferent ducts of adult mice was analyzed. Samples from five pairs of mice were used. Approximately ten videos were recorded per sample, and about seven regions of interest (ROIs) were selected for analysis from each video. A total of 432 ROIs were included for the control group, and 491 ROIs for the cKO group. The cKO mice exhibited a significant reduction in ciliary beat frequency. **** P<0.0001 compared to the control group.

To evaluate the effect of disruption of *Ift20* for cilia formation in the EDs, immunofluorescence staining was conducted using acetylated tubulin as a marker. In contrast to the robust signal clearly visible in the control group, the signal for IFT20 in the EDs of cKO mice was significantly reduced or completely absent by immunostaining (**Figure 2B, Supplementary Figure 5**). This finding indicates that presence of IFT20 is crucial for the proper formation of motile cilia in the EDs.

Histology of the EDs was examined. In control mice, few spermatozoa were present in the lumen, and long cilia extended from epithelial cells to the lumen. In the cKO mice, a striking accumulation of spermatozoa, forming a high-density mass within the ductal lumen, was evident (**Figure 2C).** The length of cilia was markedly shortened or absent in cKO mice (**Figure 2D**).

To further investigate the function of the few motile cilia observed in the EDs from cKO mice, we isolated EDs from 4-month-old control and cKO mice and analyzed their ciliary beat frequency using the Sisson-Ammons Video Analysis system (SAVA, v2.1.2w). The results revealed a significant decrease in beat frequency of motile cilia in EDs from cKO mice (**Figure 2E, Supplementary Videos 1, 2**).

### Ultrastructural defects in the motile cilia of the EDs

To further analyze the abnormal formation of motile cilia, we performed transmission electron microscopy (TEM) on collected EDs. Compared to the control group, the ciliogenesis nucleation structures (basal bodies; BB) in the cKO mice were mostly normal in ultrastructure, exhibiting the characteristic 9+0 microtubule arrangement (**Figure 3, Supplementary Figure 6A**); however, their number was significantly reduced (**Supplementary Figure 6B**). Importantly, cKO cilia were shorter than control, with some cilia ceasing extension shortly after protruding from the plasma membrane (**Figure 3**), suggesting that elongation was impaired. In cross-sectional views of cKO cilia, the typical central pair (CP) of microtubules was seldom observed, and the peripheral doublet microtubules (DMTs) were partially missing and disorganized. When few DMTs were visible, the inner and outer dynein arms of the motor complexes, were poorly defined (**Figure 3**). These findings suggest that the absence of IFT20 disrupts both early formation and elongation steps of motile cilia from EDs, and potentially the delivery of dynein motors to the ciliary axoneme.

**Figure 3.**
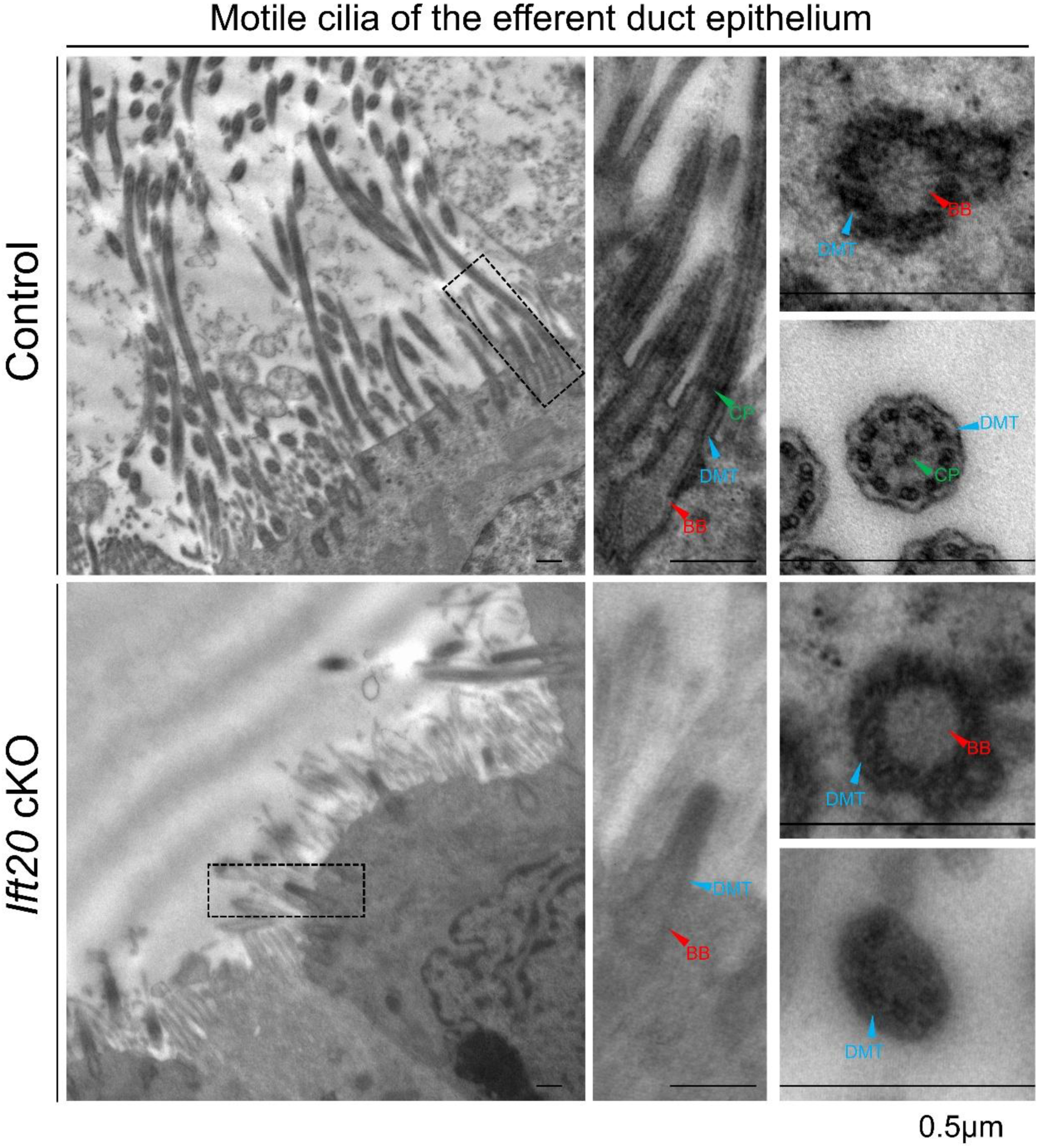
Transmission electron microscopy reveals structural defects in the motile cilia of the efferent ducts. Transmission electron microscopy analysis showed that control mice exhibited normal motile ciliary ultrastructure, featuring a 9+0 configuration at the basal body (BB) and the classic 9+2 microtubule arrangement in the axoneme section. In contrast, cKO mice displayed sparse motile cilia with impaired elongation, as the majority of cilia ceased growth shortly after initiation. Although the basal bodies retained a 9+0 structure, the axonemal microtubule organization was disrupted, with partial absence of both the outer doublet microtubules (DMTs) and the central pair (CP) microtubules. The scale is in the corner.

### Impaired protein transport in the motile cilia of EDs in *Ift20*-deficient mice

Multiple ciliary assembly-related proteins, such as SPAG16L and SPAG17, which contribute to the central apparatus of motile cilia axonemes [**35,36**], as well as DNAH2 and DNAI2, which are components of the axonemal inner and outer dynein arm complexes respectively [**37,38**], and serve as markers of dynein complex assembly in cytoplasmic DynAPs, where they are fully assembled before being subsequently transported to cilia [**39**] are essential for cilia function. We hypothesized that the transport of these proteins might be disrupted due to the absence of IFT20. Using immunofluorescence, we examined the localization of these proteins. In control mouse samples, clear fluorescent signals for DNAH2 and DNAI2 were detected both in the cytoplasm and within motile cilia. In contrast, in cKO mice, the signals were predominantly cytoplasmic and showed apparent accumulation **(Figure 4A, B)**, suggesting impaired transport of these two proteins to the cilia. SPAG16L and SPAG17 were abundantly present in the cytoplasm, with relatively weaker signals in motile cilia. This likely reflects their primary role in forming the central apparatus proteins, which may be less accessible to antibodies. Nevertheless, we could observe SPAG16L and SPAG17 signals within the motile cilia of control mice, whereas in cKO mice, their signals were primarily concentrated in the cytoplasm **(Figure 4C, D)**, with weak signals in the ciliary compartment, albeit fewer motile cilia were formed. These findings suggest that the presence of IFT20 is required for the transport of DNAH2, DNAI2, SPAG16L, and SPAG17 within the EDs epithelia, and that the failure of motile ciliogenesis is likely associated with the failure of their transport.

**Figure 4.**
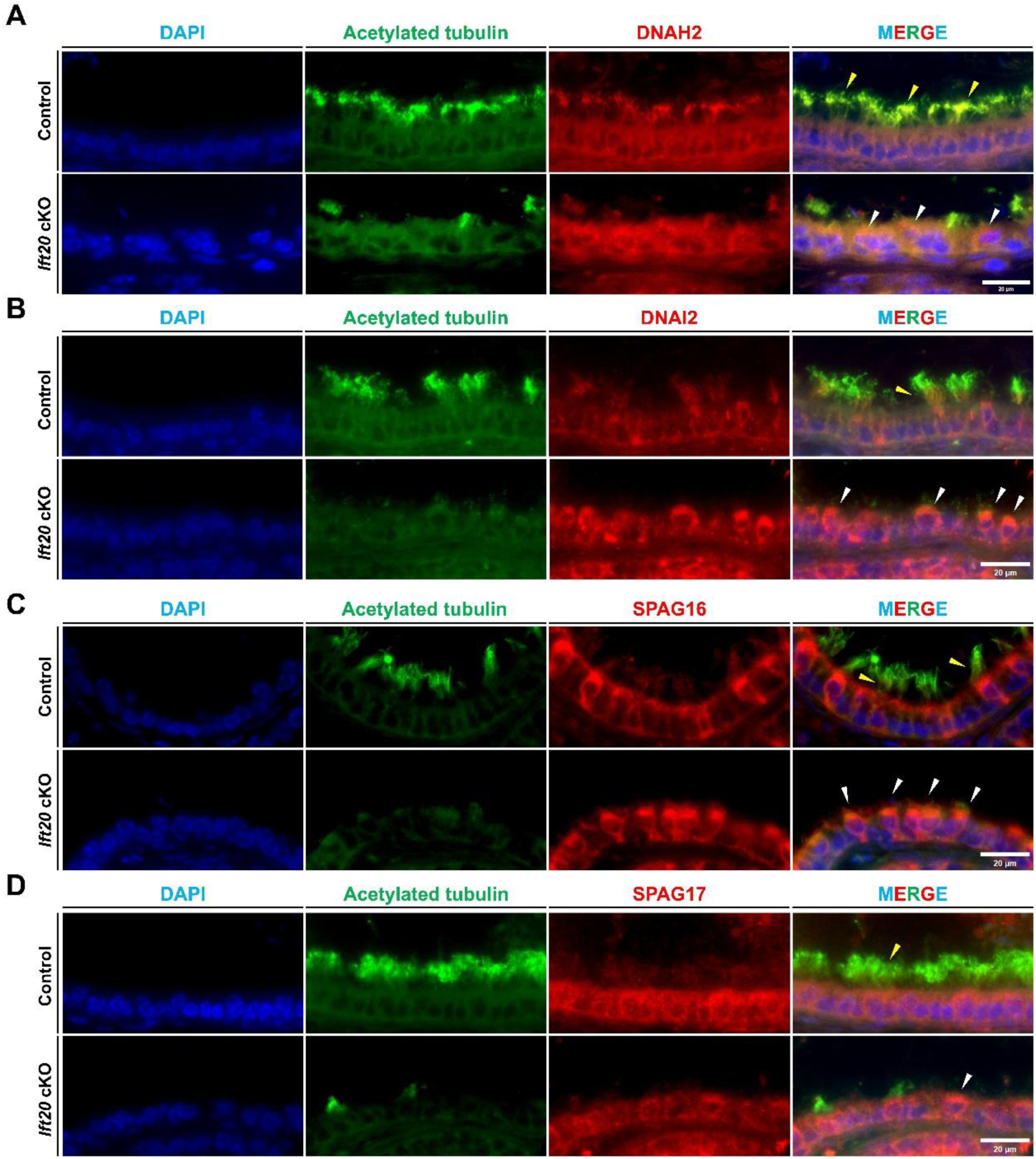
The efferent ducts of cKO mouse exhibit transport defects of motile cilia-associated proteins. Immunofluorescence staining was conducted to analyze localization of DNAH2 (**A**), DNAI2 (**B**), SPAG16 (**C**), and SPAG17 (**D**) in the efferent ducts from 4-month-old control and cKO mice. These proteins are detected in the cilia in the control mice (yellow arrows). However, strong signals were present in the apical regions inside the cytoplasm in the cKO mice (white arrowheads), indicating potential sites of protein accumulation. Green: acetylated α-tubulin, motile cilia marker; Blue: DAPI. Scale bar is shown in the corner.

### Spermatogenic defects and abnormal sperm transportation in the epididymis of *Ift20*-deficient mice

To further investigate the underlying cause of the impaired fertility in cKO mice, we analyzed histology of testes, and epididymides from the control and cKO mice. Compared to those of littermate controls, the seminiferous tubules of most cKO mice exhibited an enlarged lumen and a thinning of the epithelial layer (**Figure 5A, B, C**). Few sperm were observed inside the lumen of the seminiferous tubules (**Figure 5A, E**). Even though sperm are present throughout the epididymis in the control mice. They are rarely observed in the caput, corpus, or cauda epididymis of most cKO mice (**Figure 5A**), further suggesting that sperm transport was obstructed at the level of the EDs, preventing their normal passage.As reported[**29**], the absence of motile cilia in the EDs disrupts the maintenance of sperm in suspension, impairs fluid reabsorption, increases intraluminal pressure, and transmits this pressure back to the testis, ultimately damage the seminiferous epithelium. To evaluate seminiferous epithelium damage in cKO mice, we performed TUNEL assay on testicular sections. The results showed a significant increase in germ cell apoptosis within the seminiferous tubules adjacent to the rete testis (**Figure 5D**, **Supplementary Figure 7**). Interestingly, approximately 70% cKO mice exhibited a complete absence of sperm in the epididymis, in association with shortened/malformed motile cilia in their EDs. In some cKO mice, sperm were present in the caput, corpus, and cauda epididymis, but no obvious sperm aggregation or clumping was observed (**Supplementary Figure 8A, C**). Analysis of epididymal sperm smears from these mice revealed a significantly elevated percentage of headless sperm (**Supplementary Figure 8B**). This phenotype may potentially be associated with increased epididymal pressure. However, the testes of these mice did not show significant changes in size or seminiferous epithelial thinning, at least not by four months of age. In summary, the absence of IFT20 leads to defective formation of motile cilia in the EDs, resulting in obstructed sperm outflow. This abnormality is followed by secondary contraction of the seminiferous epithelium, accompanied by increased germ cell apoptosis.

**Figure 5.**
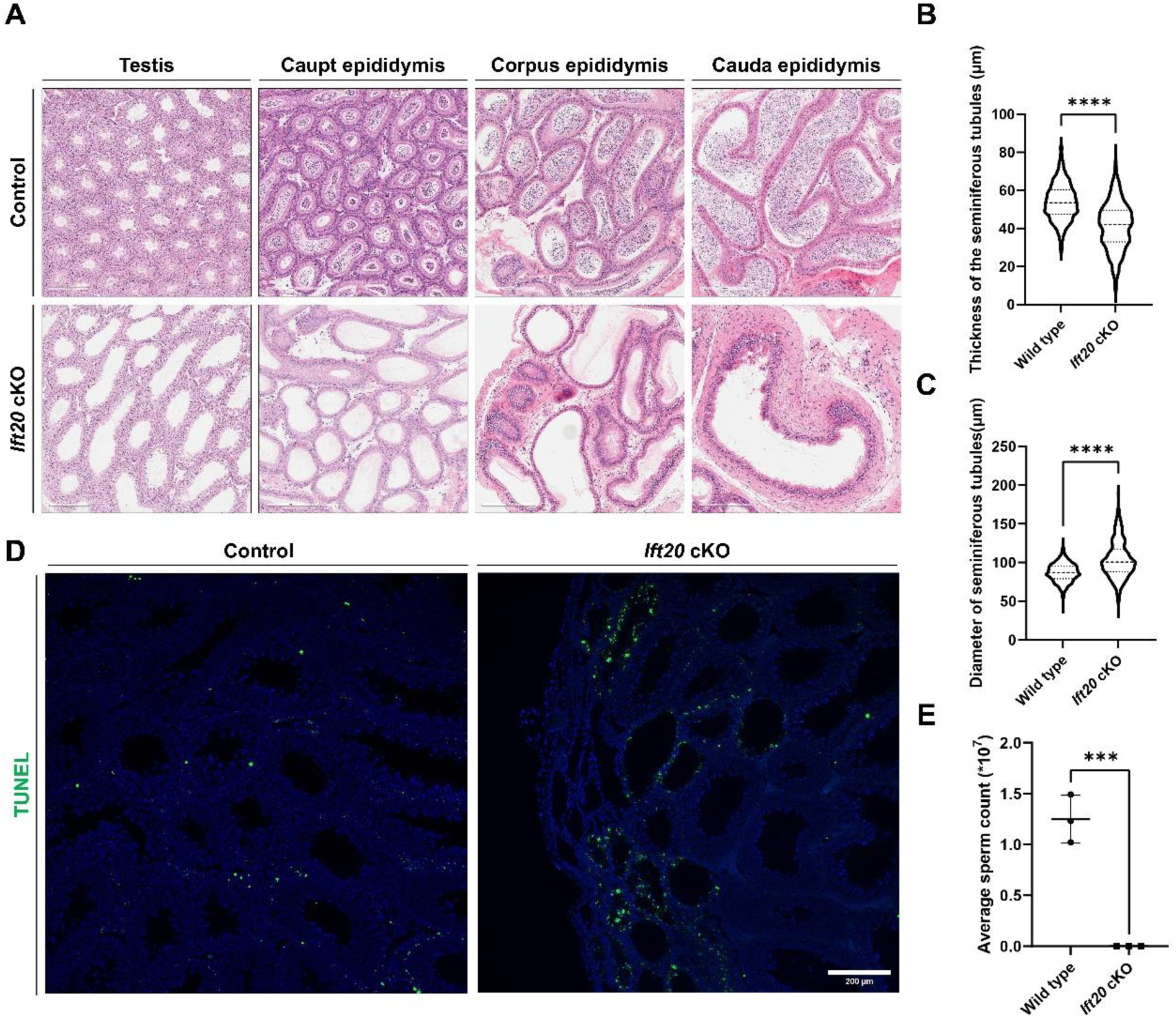
Inactivation of *Ift20* in the efferent ducts causes disruption of seminiferous epithelium followed by the absence of sperm in the epididymis. A: Tissues from 4-month-old mice were selected for HE staining. The seminiferous epithelium of cKO mouse testes showed expansion, and the epididymis was devoid of released spermatozoa in all three segments. **B**: Seminiferous tubule epithelial thickness was measured in testicular samples from three pairs of mice. A total of 1395 tubules were analyzed from control mice (mean 53.51 μm), while 1123 tubules were included from cKO mice (mean 42 μm). **** P<0.0001 compared to the control group. **C**: The diameters of seminiferous tubules were also measured using testicular samples from three pairs of mice. A total of 1473 tubules were analyzed from control mice (mean 86.97 μm), while 1071 tubules were included from cKO mice (mean 100.4 μm). **** P<0.0001 compared to the control group. **D**: Apoptosis in the testis was examined using 4-month-old mice. While minimal green apoptotic signals were detected in the testes of control mice, extensive apoptotic signals were observed near the rete testis in cKO mice. **E**: Sperm counts from the cauda epididymis were performed. The cauda epididymis was dissected from the mice, minced, and incubated in 1 ml of pre-warmed PBS at 37°C for 10 min. Sperm were subsequently counted using a standard hemocytometer. A significant reduction in sperm count was observed in cKO mice that showed typical seminiferous duct dilation phenotype (n=3). *** P<0.001 compared to the control group.

### Short motile cilia and reduced beat frequency in the tracheal epithelium of *Ift20*-deficient mice

Our previous study on IFT140 conditional knockout mice revealed severe defects in ciliogenesis in the trachea and intrapulmonary airways [**30**]. In contrast, almost all IFT20-deficient cKO mice survived to adulthood. To investigate whether the motile cilia of the trachea in these cKO mice exhibited similar phenotypic abnormalities, and after confirming efficient deletion of IFT20 (**Figure 6A**), we collected tracheas from cKO mice for histological analysis. Hematoxylin-Eosin staining indicated that approximately one-third of the cKO mice exhibited shortened and relatively sparse tracheal motile cilia (**Figure 6B, C**). However, in the remaining cKO mice, the tracheal cilia length appeared similar to controls (**Supplementary Figure 9**). Interestingly, despite this variability in length, ciliary beat frequency was significantly reduced across all cKO mice (**Figure 6D**, **Supplementary Videos 3, 4**).

**Figure 6.**
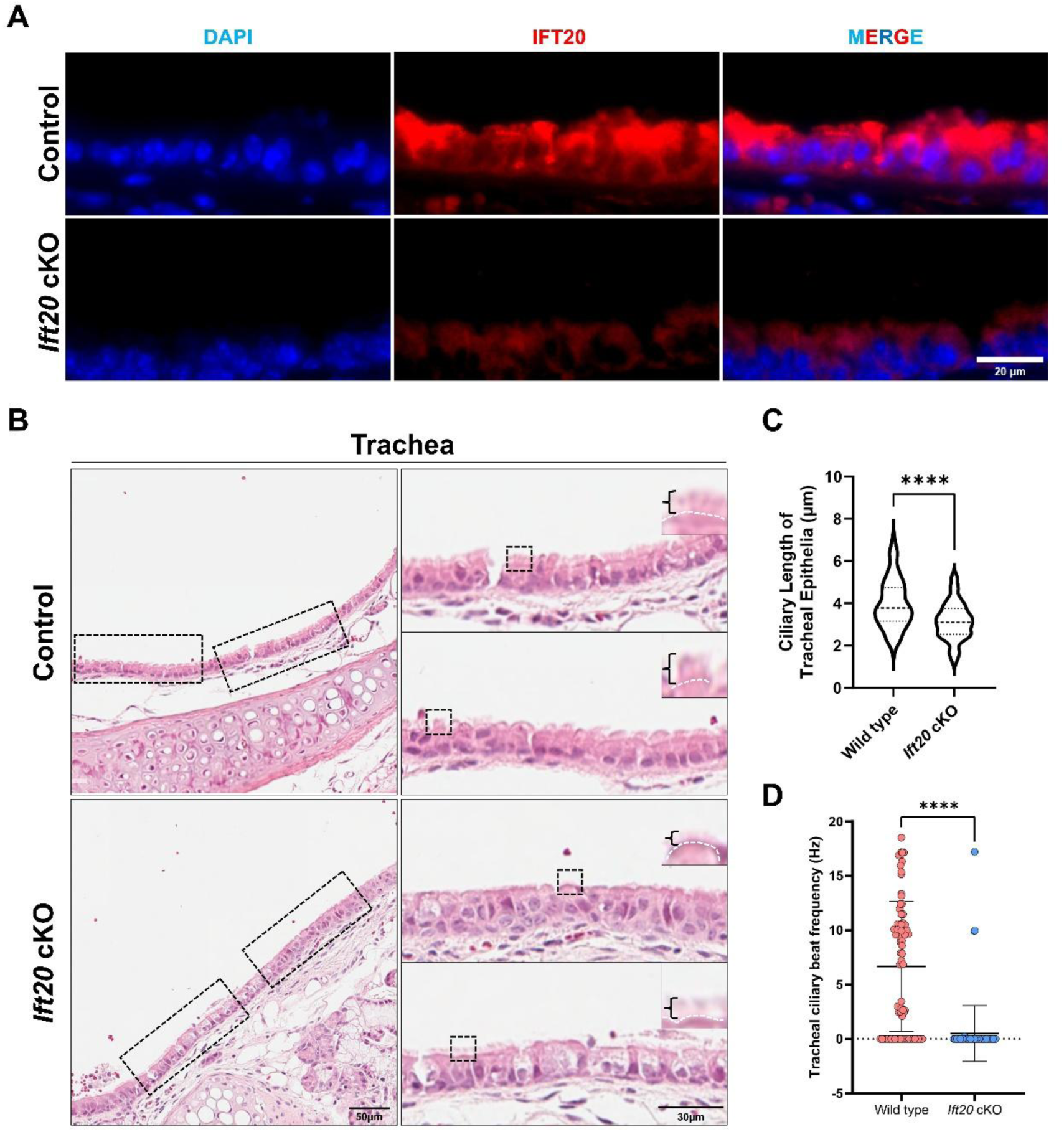
Loss of IFT20 in the tracheal epithelium results in shortened motile cilia length and reduced beating frequency. **A**: Representative immunofluorescence results showed that IFT20 was completely ablated in the tracheal epithelium of cKO mice. **B**: Representative HE staining results revealed shortened motile cilia in the tracheal epithelium of 4-month-old cKO mice. **C**: Cilia length measured from histology images. Based on HE staining, 4 pairs of mice were selected, and 83 regions of interest (ROIs) each from the control and cKO groups were measured. The results showed that, compared to the control group (mean 3.985 µm), cilia length was shortened in cKO mice (mean 3.152 µm). **D**: Reduced ciliary beating frequency in the cKO mice. Using the same video recording method as for the efferent duct motile cilia, videos of tracheal epithelial motile cilia were captured. Five pairs of mice were selected for statistical analysis. A total of 130 ROIs were included in the control group (mean 3.089 Hz), and 82 ROIs were included in the cKO group (mean 0.3645 Hz). Notably, during video analysis, we observed that unlike in the efferent ducts, the tracheal tissue is thicker due to the presence of cartilage, and more fluid is trapped above the motile cilia. This likely caused the analysis software to filter out more beating data as background noise. Therefore, these results represent relative values after background noise removal, which may explain why a significant portion of the data showed a beating frequency of 0 Hz.

### Impaired protein localization in tracheal epithelial motile cilia of *Ift20*-deficient mice

We also examined the trachea by transmission electron microscopy. In the vast majority of cKO mice, the motile cilia appeared structurally normal, possessing the typical “9+2” microtubule arrangement (**Figure 7**). The characteristic obstructions observed in the IFT140 knockout model, either along the ciliary shaft or at the ciliary tip, were not evident, likely reflecting the distinct roles of IFT20 and IFT140 within the IFT system. Subsequently, we performed immunofluorescence staining to localize cilia structures. While outer and inner dynein arm complexes marked by DNAH2 and DNAI2 respectively, appeared to be transported to the motile cilia in the tracheal epithelium of cKO mice, cytoplasmic accumulation of these proteins was also observed, suggesting either partial impairment of protein transport (or an accumulation of ciliary structures destine for cilia that did not achieve full length) (**Figure 8A, B**). The presence of at least some outer and inner dynein arm proteins in cilia also suggests that the effects of IFT20 loss may be partially compensated for by alternative pathways. SPAG16 and most of the SPAG17 signals appeared to be normally localized to the motile cilia, with comparable signals between cKO and control tracheal epithelium (**Figure 8C**). However, in one cKO mice (n=6), SPAG17 failed to localize to the motile cilia, with its signal remaining at the ciliary base and within the cytoplasm (**Supplementary Figure 10**). In summary, the findings suggest that IFT20 deficiency disrupts the transport of cilia proteins, cilia assembly, and subsequently normal cilia beating.

**Figure 7.**
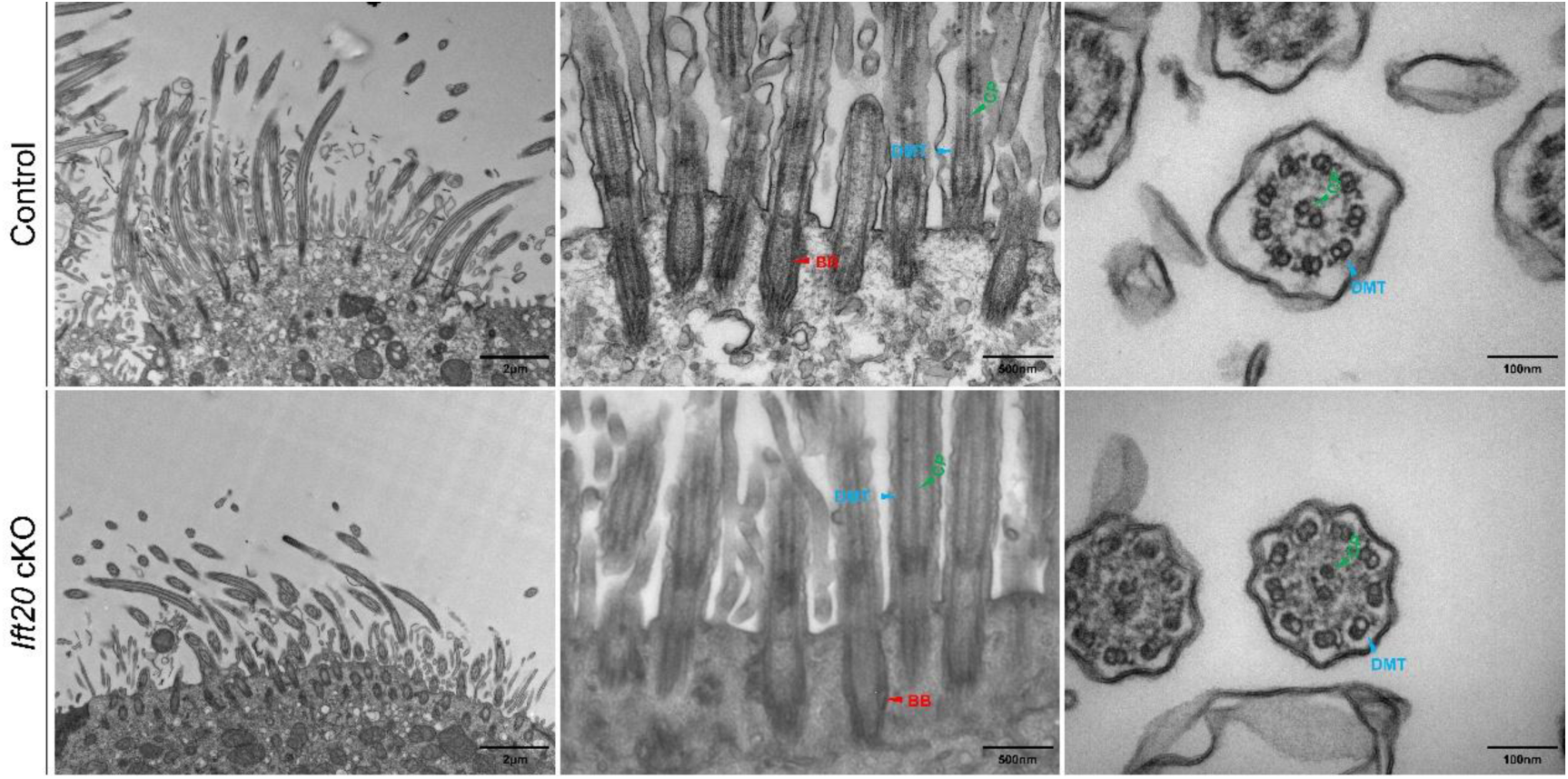
Transmission electron microscopy showed no significant changes in the ultrastructure of the tracheal epithelium of cKO mice. Transmission electron microscopy of tracheal tissues from 4-month-old mice revealed that both cKO and control groups exhibited typical motile cilia ultrastructure, including basal bodies anchored beneath the plasma membrane and the classic 9+2 microtubule arrangement in the axoneme. No obvious abnormalities were observed in the inner or outer dynein arms.

**Figure 8.**
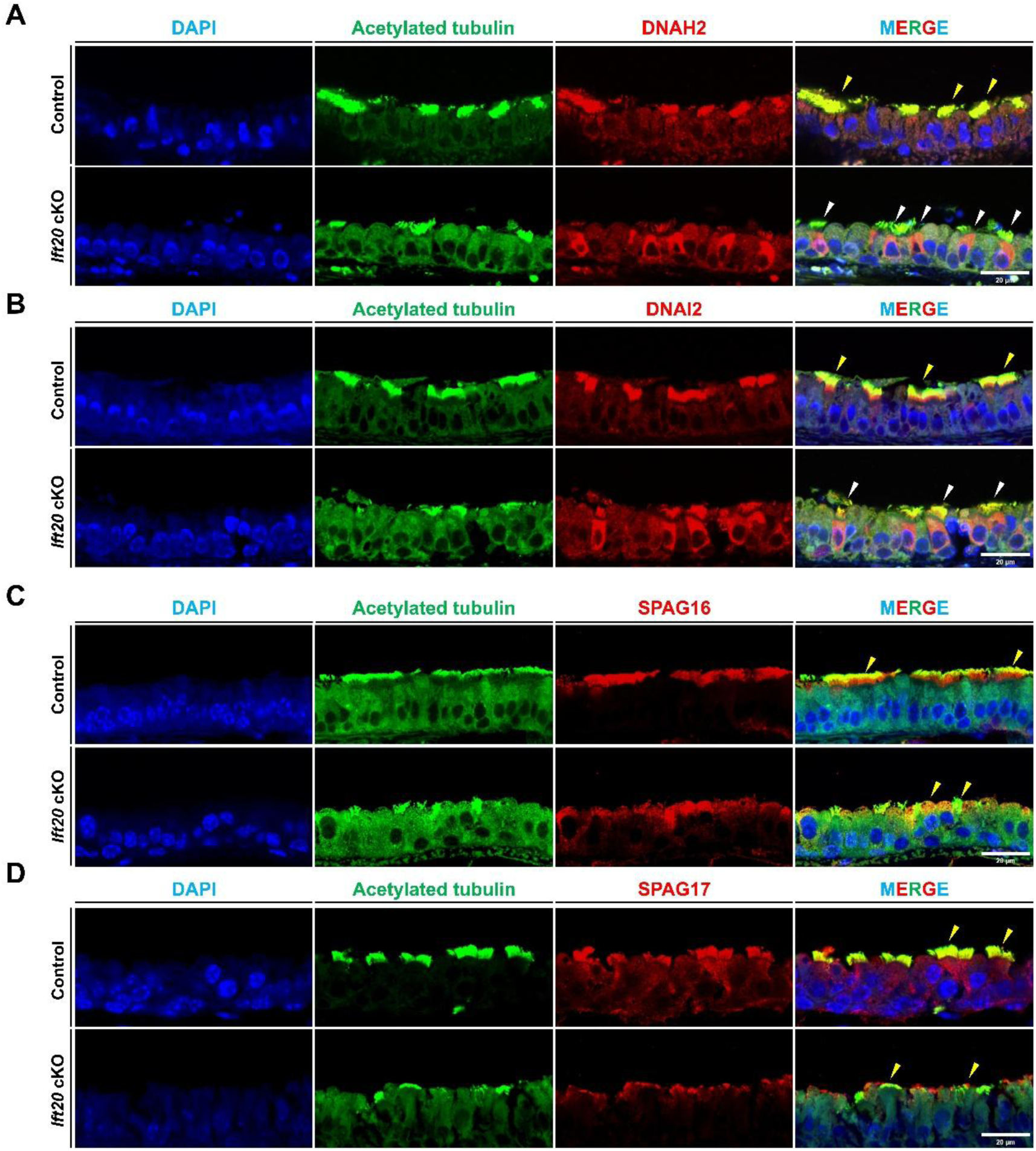
The trachea of cKO mouse exhibit transport defects of motile cilia-associated proteins. Immunofluorescence was performed to assess the expression and localization of DNAH2(**A**), DNAI2(**B**), SPAG16(**C**), and SPAG17(**D**). These proteins are detected in the cilia in the control mice (yellow arrows). White arrows indicate potential sites of protein accumulation. Green: acetylated α-tubulin, motile cilia marker; Red: target protein; Blue: DAPI. Scale bar is shown in the corner.

### Study limitations

Several limitations of this study should be acknowledged. First, although FOXJ1-Cre drives recombination in most motile ciliated cells, we only examined efferent ductules and trachea. The role of IFT20 in other motile ciliated organs, such as brain ependyma and oviducts, was not systematically evaluated; notably, female fertility data were insufficient to draw firm conclusions. Second, the milder and heterogeneous tracheal phenotype could not be fully explained, and potential compensatory mechanisms or variable Cre recombination efficiency were not further investigated. Third, the evidence for impaired protein transport relies primarily on immunofluorescence localization; direct assessments of intraflagellar transport dynamics, cargo trafficking kinetics, or rescue experiments are lacking. Finally, ultrastructural defects were described qualitatively without quantitative analysis of their incidence. Addressing these limitations in future studies will help further clarify the tissue-specific functions of IFT20 in motile cilia.

## Discussion

Here, we demonstrate a critical and tissue-dependent role for the intraflagellar transport (IFT) protein IFT20 in the formation and function of mammalian motile cilia. Since the initial discovery of intraflagellar transport (IFT) in the flagella of *Chlamydomonas* by Kozminski [**40**], IFT has been extensively studied in diverse organisms, and its underlying mechanism is relatively conserved [**4**]. In unicellular organisms such as the green alga *Chlamydomonas reinhardtii*, IFT mediates the transport of ion channel proteins into the cilium, where they participate in light perception and regulate flagellar beating [**41–44**]. IFT20, along with other IFT components, was first identified in *Chlamydomonas reinhardtii* in 1998 [**45**]. Subsequently, its localization pattern at the primary cilium and the ciliary basal body was characterized in mammalian photoreceptor cells [**46**]. Later, John *et al*. demonstrated that IFT20 also prominently localizes to the Golgi complex and shuttles between the Golgi and the primary cilium [**15**], suggesting its involvement in the transport of proteins from the Golgi to the primary cilium. Subsequent research has further elucidated the diverse roles of IFT20 in primary cilia biology [**25**].However, it remained unclear whether IFT20 participates in the formation and assembly of motile cilia in multiciliated cells, although our previous study has demonstrated that IFT20 is essential for the assembly of sperm flagella [**27**]. Here, we addressed this gap of knowledge by characterizing the phenotypic consequences of IFT20 loss in two motile cilia-rich tissues: the EDs and the trachea.

### IFT20 deficiency results in progressive and tissue-specific motile cilia defects

IFT20 showed similar expression levels in the lung and EDs, with both being significantly lower than in the testis, while its expression in the brain was lower than all three tissues. Our study reveals a stark contrast in the phenotypic severity between these tissues. In the EDs, IFT20 loss led to a severe, early-stage ciliogenesis defect, characterized by a marked reduction in basal body number and extremely short cilia. In the trachea, however, the phenotype was significantly milder, with only a slight shortening of cilia and a reduction in CBF, while most mice survived normally. Notably, and consistent with our previous IFT140 knockout model[30], we did not observe obvious hydrocephalus in IFT20-deficient mice. While ependymal cilia dysfunction is often linked to hydrocephalus, this finding adds to the ongoing debate about its causality [**47**] and further underscores the remarkable tissue-specificity of IFT requirements. This tissue-specific requirement for IFT20 likely from two factors. First, intrinsic differences in the structural and functional demands of the cilia themselves may dictate their reliance on IFT-mediated maintenance. EDs cilia are longer and beat more rapidly to generate turbulent flow[**29**], potentially creating higher energy and turnover demands that make them more susceptible to IFT disruption. In contrast, airway cilia are shorter, beat more slowly and coordinately [**48**], and may be less dependent on constant, high-flux IFT. Second, and importantly, we observed a significant reduction in basal body number in the EDs, suggesting that IFT20 is not only involved in ciliary maintenance but also in the stability or generation of basal bodies themselves in this tissue. However, due to experimental limitations, the specific stages at which IFT20 participates in this regulation remain to be elucidated.

### IFT20 differentially affects the transport of specific ciliary protein complexes

Analysis of ciliary protein transport revealed that IFT20 deficiency impairs the trafficking of specific cargoes, and this effect is also tissue-dependent. In both tissues, the loss of IFT20 disrupted the transport of markers dynein arm complexes, DNAH2 and DNAI2, which accumulated at the ciliary base and in the cytoplasm. However, some of these proteins still managed to localize within the remaining cilia, implying the existence of compensatory mechanisms that partially mitigate the loss of IFT20. More striking was the differential effect on central apparatus proteins. While SPAG16L appeared to be transported normally to tracheal cilia, SPAG17 failed to localize to cilia in a subset of tracheal epithelial cells, remaining trapped in the cytoplasm **(Supplementary Figure 10)**. This suggests that IFT20 may have distinct transport capacities for different central apparatus components. In contrast, the severe ciliogenesis failure in the EDs made it difficult to assess the transport of SPAG16L and SPAG17, as their signals were either absent from the rudimentary cilia or predominantly cytoplasmic, a finding that aligns with the near-complete loss of ciliary structure. This data, taken together, indicates that the primary defect in the EDs is an early block in ciliary assembly, whereas in the trachea, IFT20 is more critical for the fine-tuning of ciliary function by facilitating the transport of specific dynein motors.

### Phenotypic comparison of IFT20 and IFT140 knockout models reveals complex tissue dependencies

The tissue-specific phenotype we observe for IFT20 is further highlighted by comparing it to our previous work on IFT140, a core component of the retrograde IFT-A complex[30]. While both proteins are essential for ED motile cilia **(Supplementary Figure 11)**, but their roles in the trachea differ markedly. IFT140 deficiency causes severe, early postnatal lethality with grossly malformed cilia displaying bulbous tips, indicative of impaired IFT and cargo accumulation [40], whereas IFT20 deficiency results in a much milder, primarily functional defect. This finding is somewhat counterintuitive, as it contrasts with studies in photoreceptors [**49**] and kidney primary cilia [**31**], and suggests that the tracheal IFT machinery is not uniformly dependent on all its components. We propose two, non-mutually exclusive explanations for this discrepancy. First, other proteins or pathways may partially compensate for the loss of the small IFT20 protein in the trachea, enabling the core anterograde IFT machinery to function, albeit with reduced efficiency for specific cargoes like SPAG17. Second, and more compellingly, the distinct structural roles of IFT20 and IFT140 likely contribute to the phenotypic differences. IFT140 is a large scaffold protein critical for the structural integrity of the IFT-A complex and retrograde transport. IFT20, in contrast, is a small, peripheral component of the IFT-B complex with a known, additional role in Golgi-to-cilia trafficking [**15**]. Its absence may primarily disrupt a specific subset of cargo delivery pathways originating from the Golgi. Recent research supports the existence of multiple, parallel routes for protein delivery to the cilium [**50**]; for instance, the transport of some proteins depends on IFT20 and the exocyst complex, while others, like Smoothened, do not. We hypothesize that in the trachea, motile cilia assembly relies more heavily on IFT20-independent trafficking pathways, whereas ED ciliogenesis is critically dependent on the specific IFT20-mediated route from the Golgi. The severe ED phenotype we observe is therefore likely a combined result of impaired core IFT and a failure in the early, IFT20-dependent delivery of essential building blocks from the Golgi to the ciliary base.

### Location-Dependent Testicular and Epididymal Manifestations in Motile Cilia-Deficient Mice

Interestingly, although motile cilia defects are present in the EDs of cKO mice, the macroscopic phenotypes in the testis and epididymis are dependent on the location of sperm blockage. In the majority of mice at 4 months of age, no overt testicular atrophy was observed; however, distinct sperm agglutination was evident in the epididymis (**Supplementary Figure 8**). In contrast, in a minority of mice where sperm blockage occurred in the efferent ductules, the testes exhibited increased apoptosis and seminiferous tubule atrophy, accompanied by the absence of sperm in the epididymis (**Figures 2, 5**). We propose that these macroscopic manifestations correlate with the site of sperm blockage. When sperm are blocked in the EDs, pressure is rapidly transmitted to the testis, impairing normal spermatogenesis. Conversely, when sperm are blocked in the distal epididymis, this process is considerably slower; at least by 4 months of age, the testis remains unaffected. The unpredictability of the sperm blockage site resulting from motile cilia defects in the testis also indirectly indicates that the beating of motile cilia in the EDs is merely one of the objective prerequisites for normal sperm outflow. Previous studies [**29**] have demonstrated that the primary function of motile cilia beating is to maintain sperm in suspension and increase the contact area between the fluid and epithelial cells, thereby facilitating water reabsorption. The direct driving force for sperm outflow may originate from both the beating of motile cilia and the rhythmic contractions of the seminiferous tubules [**51,52**]; however, the relative contribution of cilia “beating” remains unknown. In summary, loss of IFT20 results in ciliary dysfunction, yet the macroscopic alterations may arise from the interplay of multiple factors.

In conclusion, our study establishes that IFT20 is crucial for mammalian motile ciliogenesis and exhibits marked tissue dependent functions. Future research will focus on elucidating the differential regulatory roles of various IFT protein subunits across different tissues. This will deepen our understanding of IFT proteins in motile cilia and provide new insights into the molecular mechanisms and potential therapeutic strategies for diseases caused by motile cilia dysfunction.

## Acknowledgements

This research was supported by Wayne State University Start-up fund, NIH grants HD105944 (ZZ), HD107579 (ZZ), and HD114311 (ZZ). Steven L. Brody is supported by NIH HL128370 and the Barnes-Jewish Hospital Foundation. Aminata Touré is funded by INSERM, CNRS, Université Grenoble Alpes and the Agence Nationale pour la Recherche (ANR-17-CE13-0023 DIVERCIL, ANR-19-CE17-0014 FLAGELOME). Gregory J. Pazour is supported by NIH grant GM060992.

## Author contributions

T. Li, A. Schmitt, J. Pan, C. Zheng, W. Li, S. Parveen, A. Yadav, M. Abdulghani, and K. Bennett performed research; G. Pazour, R. Hess, and C. Kelly contributed reagents; DH Huang, S. Brody, A. Touré and Z Zhang wrote the paper.

## Data availability

The data is available from the corresponding author upon reasonable request.

---

**Supplementary Figure 1.**
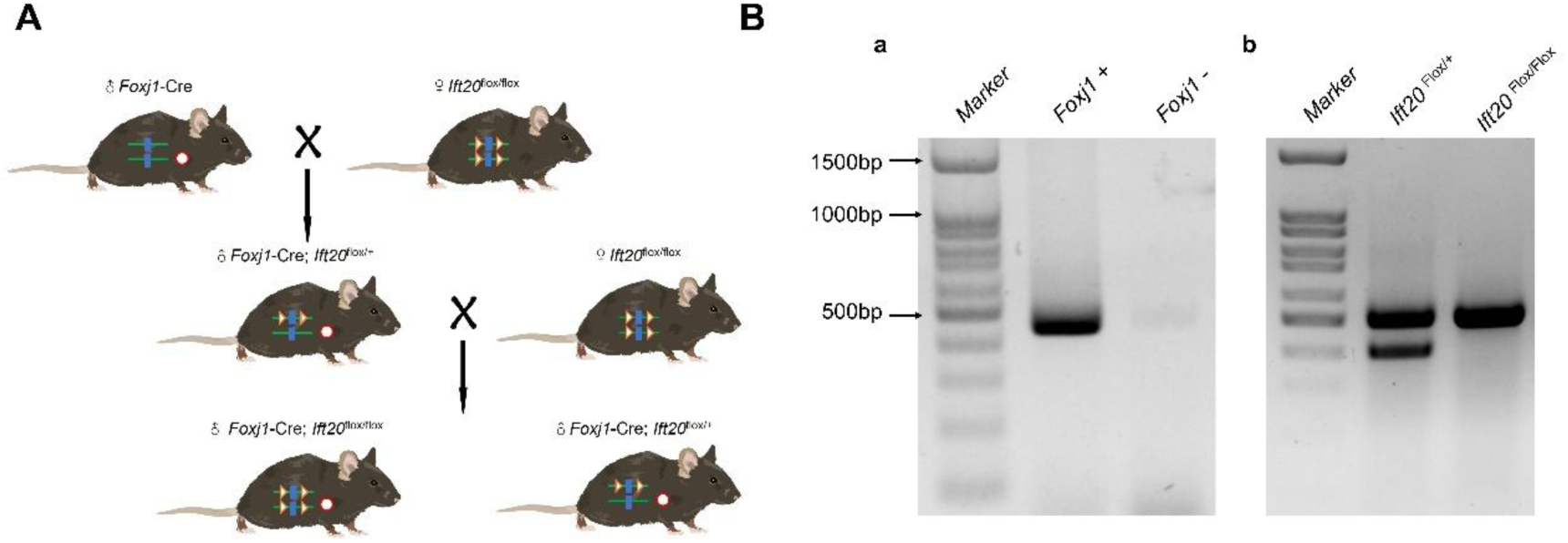
Generation of a conditional knockout mice to disrupt Ift20 in the motile cilia. A. Breeding strategy to generate the conditional knockout mice. B. A representative PCR to identify the mouse genotypes. **a.** PCR to detect the *FOXJ1*-Cre allele. **b.** PCR to detect the floxed *Ift20* allele.

**Supplementary Figure 2.**
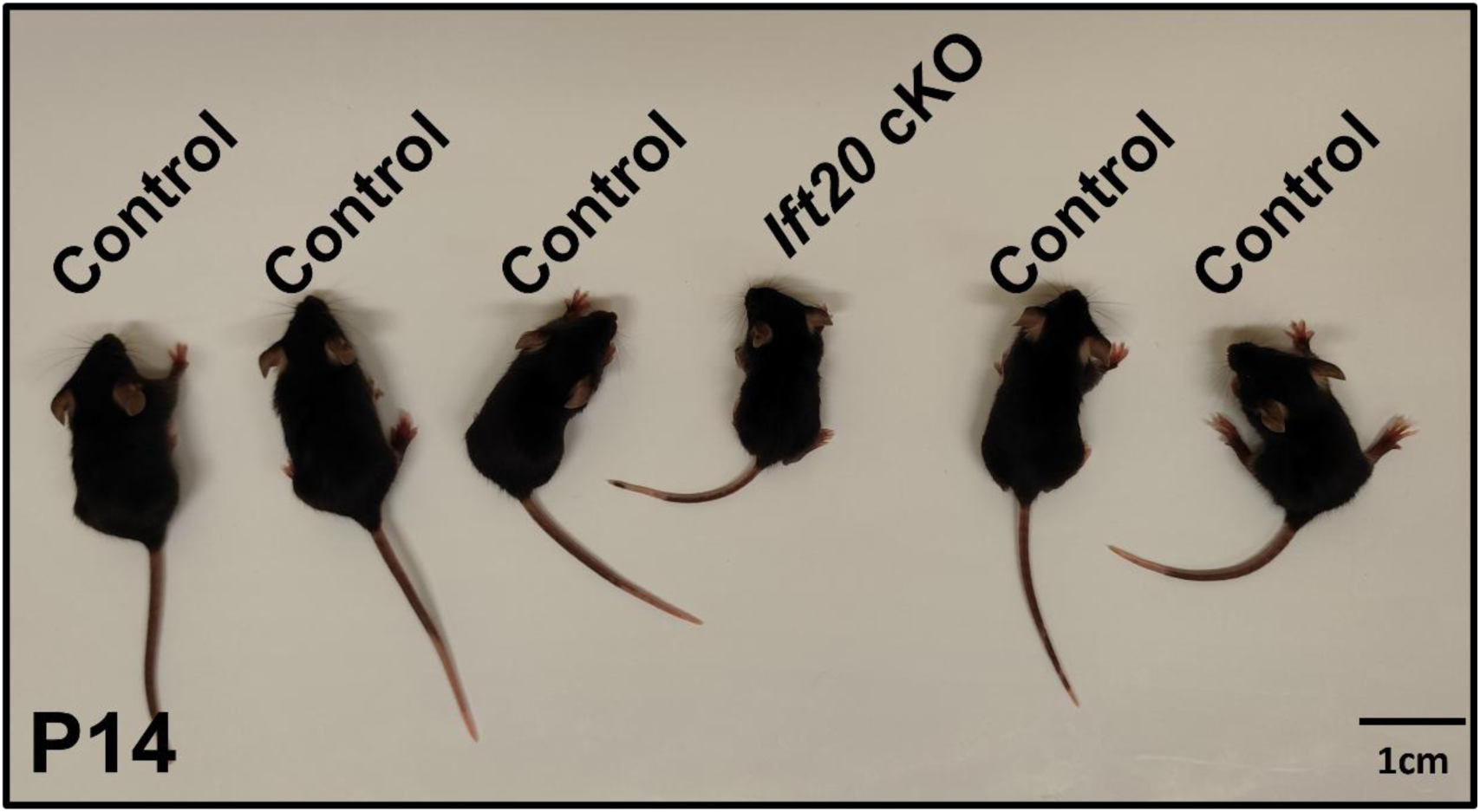
A representation image showing smaller size of 14-day-old cKO mouse. Compared to the littermates, the cKO mouse is smaller in size.

**Supplementary Figure 3.**
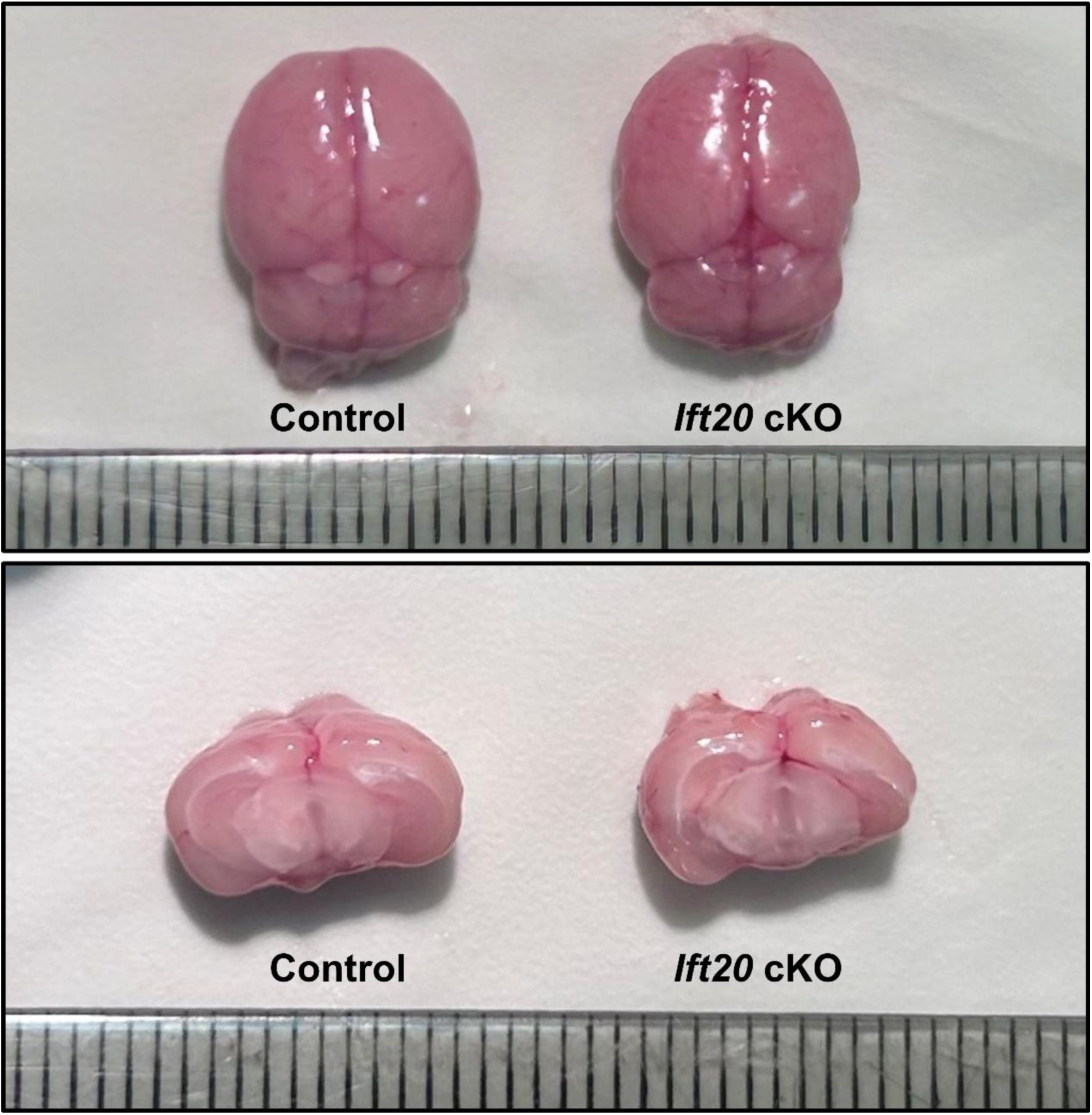
Gross examination of brain of the adult control and cKO mice. No hydrocephalus was found in the cKO mice.

**Supplementary Figure 4.**
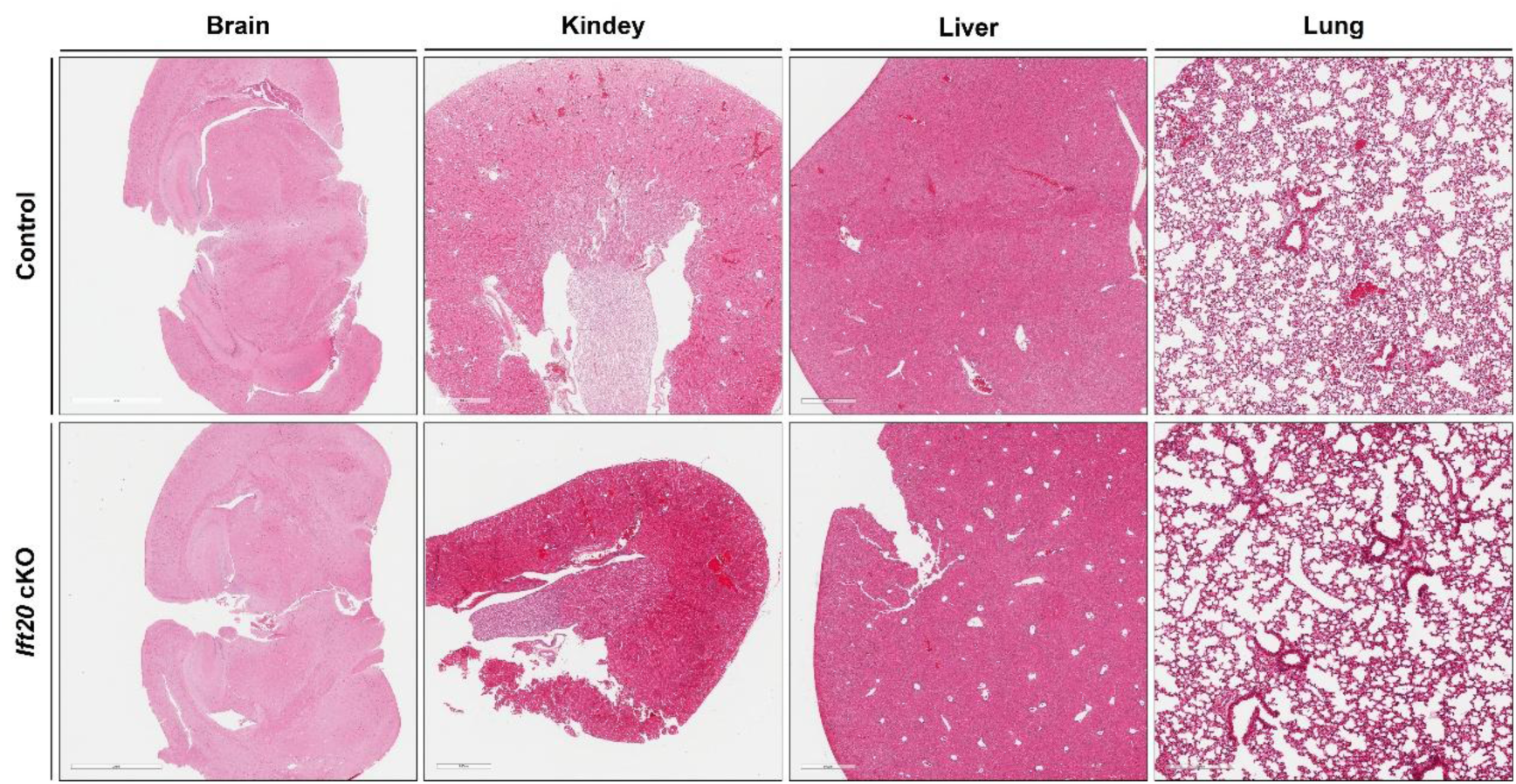
H&E staining of other organs with motile cilia after IFT20 deletion. No obvious histological changes were detected in the brain, kidney, liver, and lung tissues of cKO mice.

**Supplementary Figure 5.**
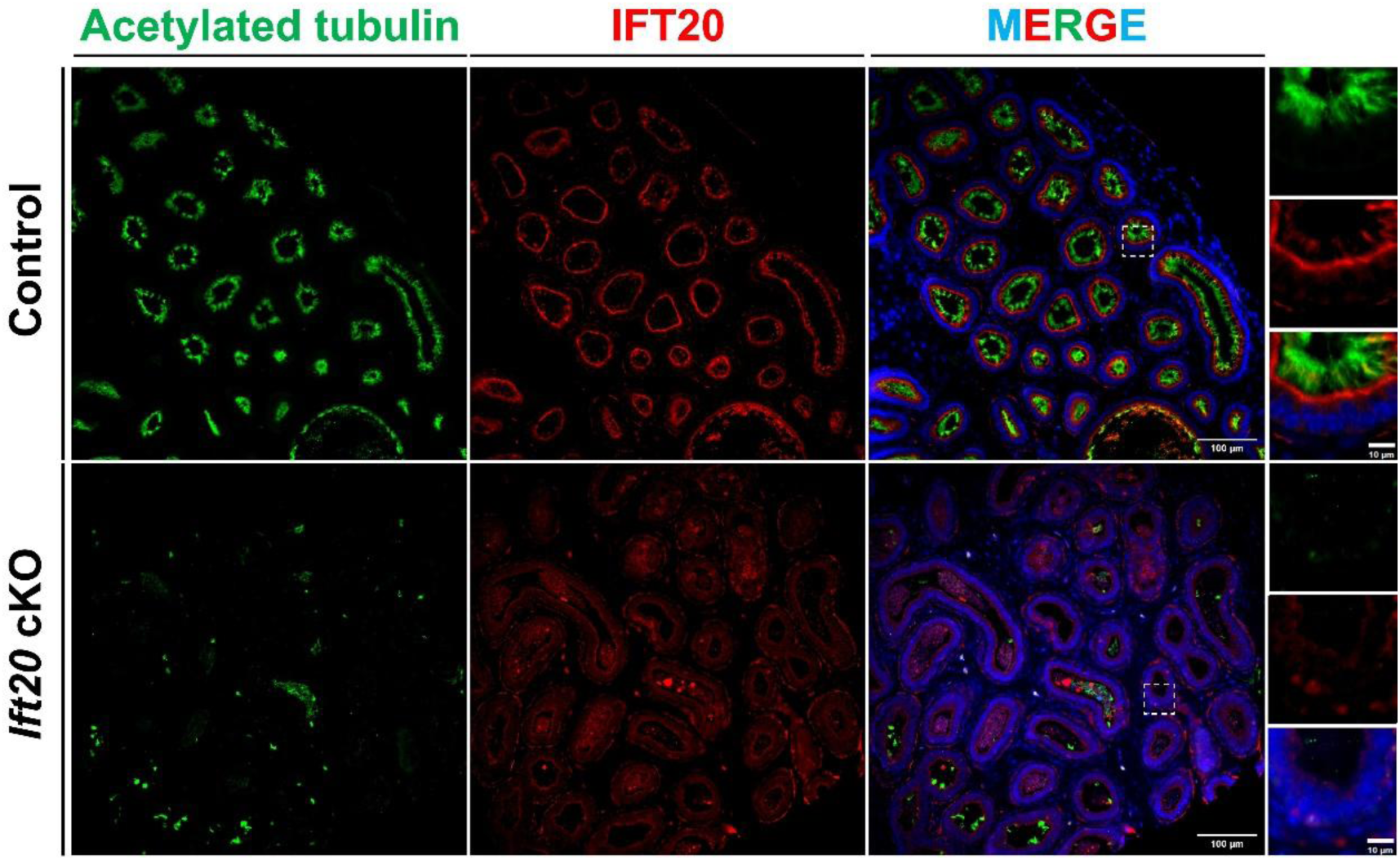
Low-intensity immunofluorescence staining results. Green: acetylated tubulin; red: IFT20; blue: DAPI. Scale bar in the corner.

**Supplementary Figure 6.**
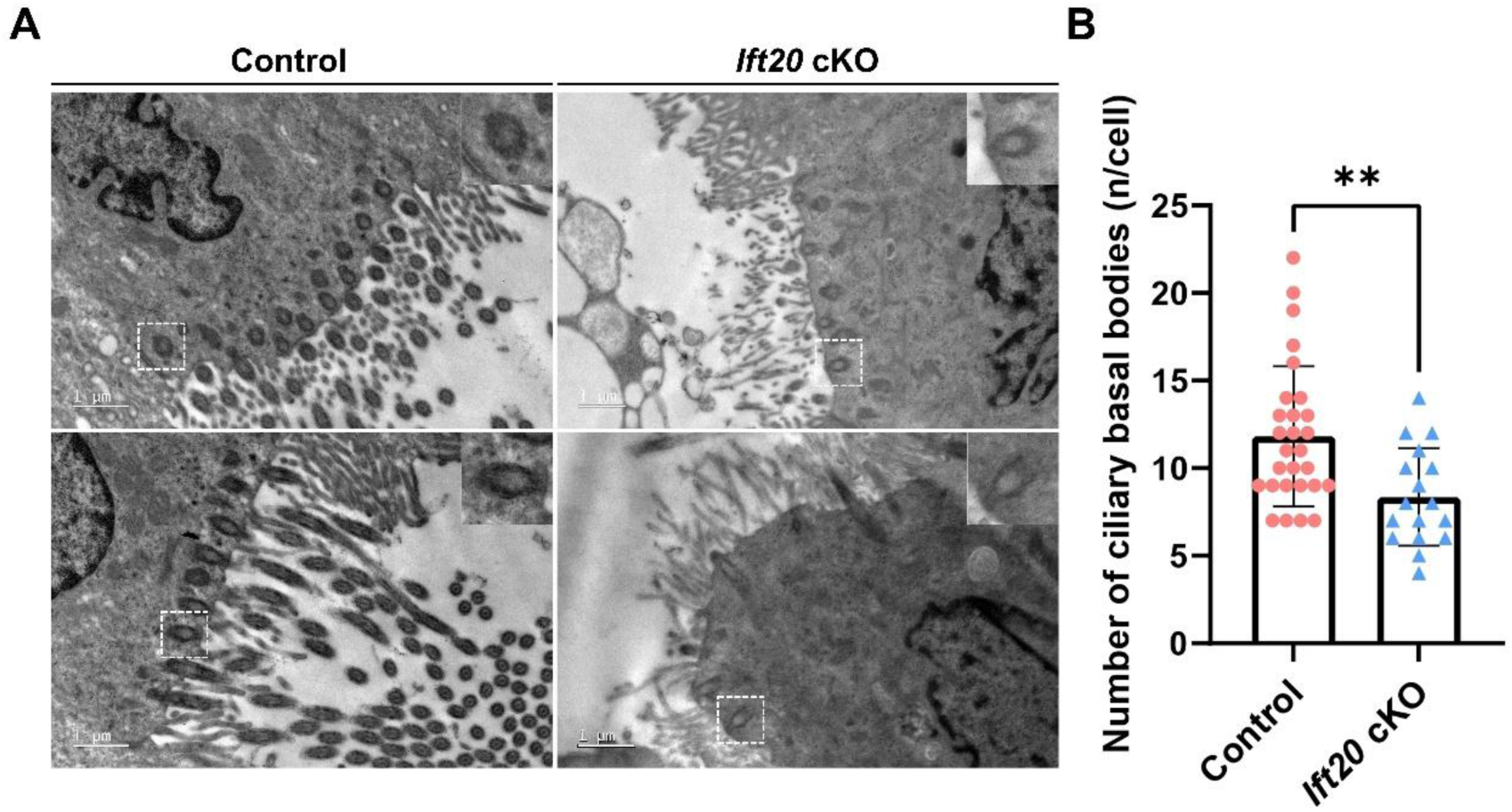
Transmission electron microscopy revealed a reduction in the number of motile cilia basal bodies in the efferent ducts of cKO mice. **A**: Transmission electron microscopy results show that epithelial cells in the control group are rich in motile cilia basal bodies, whereas the number of basal bodies is reduced in cKO mice. **B**: Based on transmission electron microscopy results, basal bodies within individual cells were counted. The control group included 28 cells (mean 11), and the cKO group included 17 cells (mean 8).

**Supplementary Figure 7.**
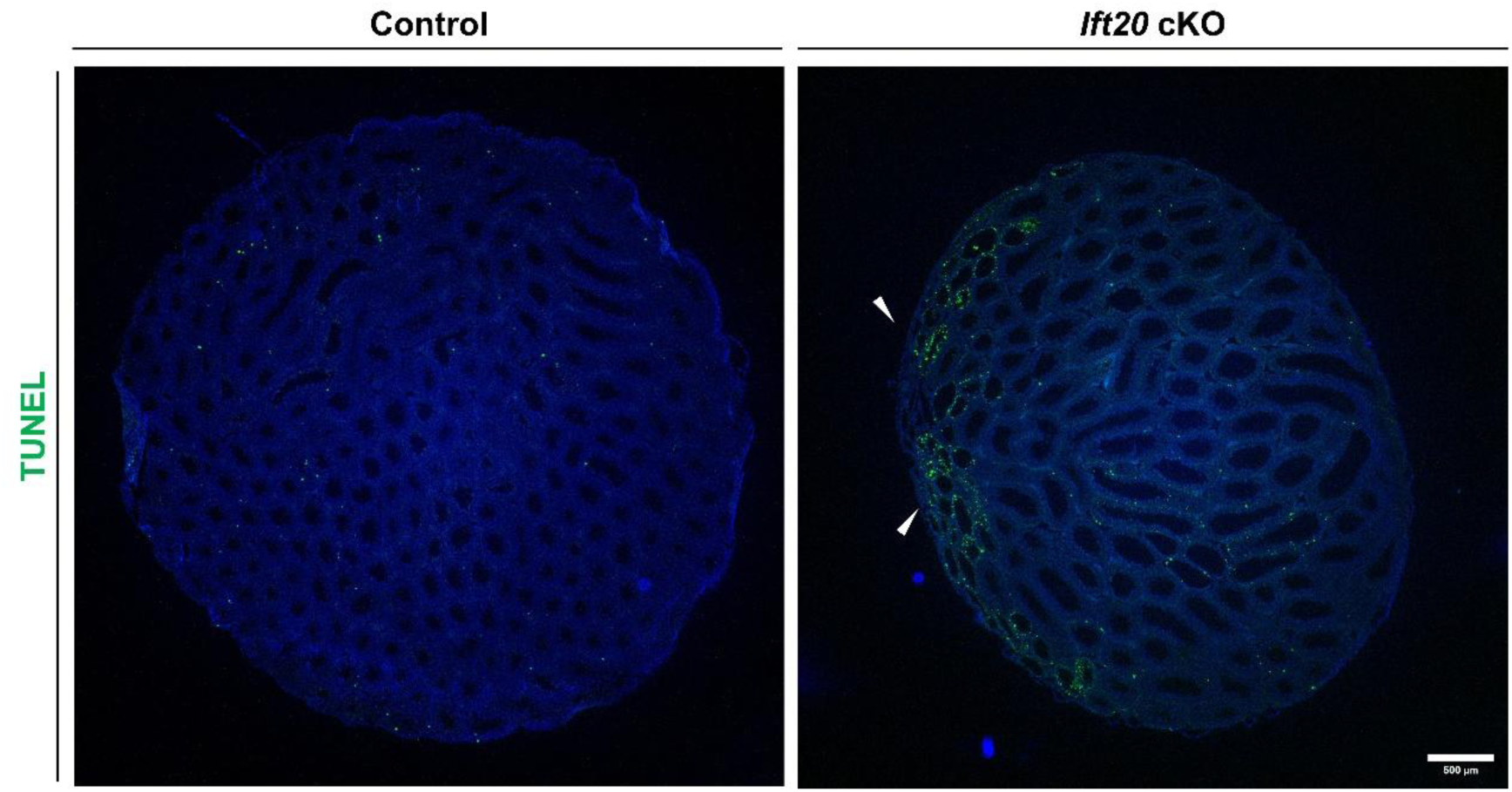
Low-magnification apoptosis immunofluorescence results. Testis samples from 4-month-old mice were selected: green apoptotic signals were rarely detected in the control group; whereas in cKO mice, apoptotic signals were primarily localized adjacent to the rete testis (white arrow). This suggests that increased ductal pressure due to impaired sperm outflow leads to cellular apoptosis.

**Supplementary Figure 8.**
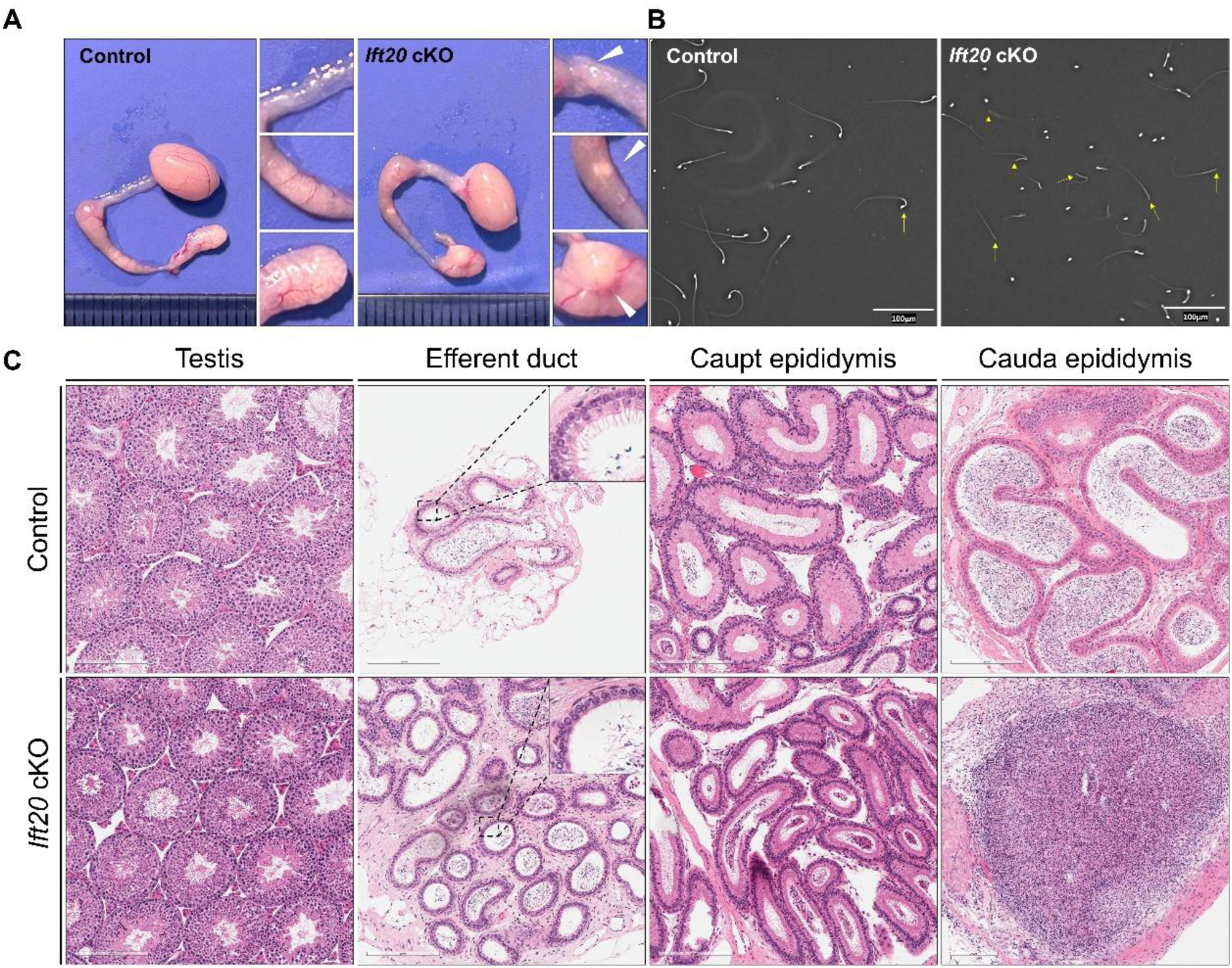
Sperm were not obstructed in the efferent ducts but rather blocked within the epididymis in some cKO mice. **A**: Photographs of the testis, efferent ducts, and epididymis showed that cKO mice exhibited distinct, white obstructive masses in the epididymis, particularly in the cauda epididymis, where hard, swollen masses were observed (white arrow). **B**: One side of the epididymis was collected, placed in PBS, minced, and incubated at 37°C for 10 min to allow sperm to swim out. After smear examination, a significant increase in the proportion of headless sperm was observed in cKO mice. **C**: HE staining revealed no swelling of the seminiferous epithelium in the testes of cKO mice. Sperm obstruction was present in both the epididymis and efferent ducts. Interestingly, due to severe obstruction in the cauda epididymis, there was a greatly enlarged lumen completely filled with sperm.

**Supplementary Figure 9.**
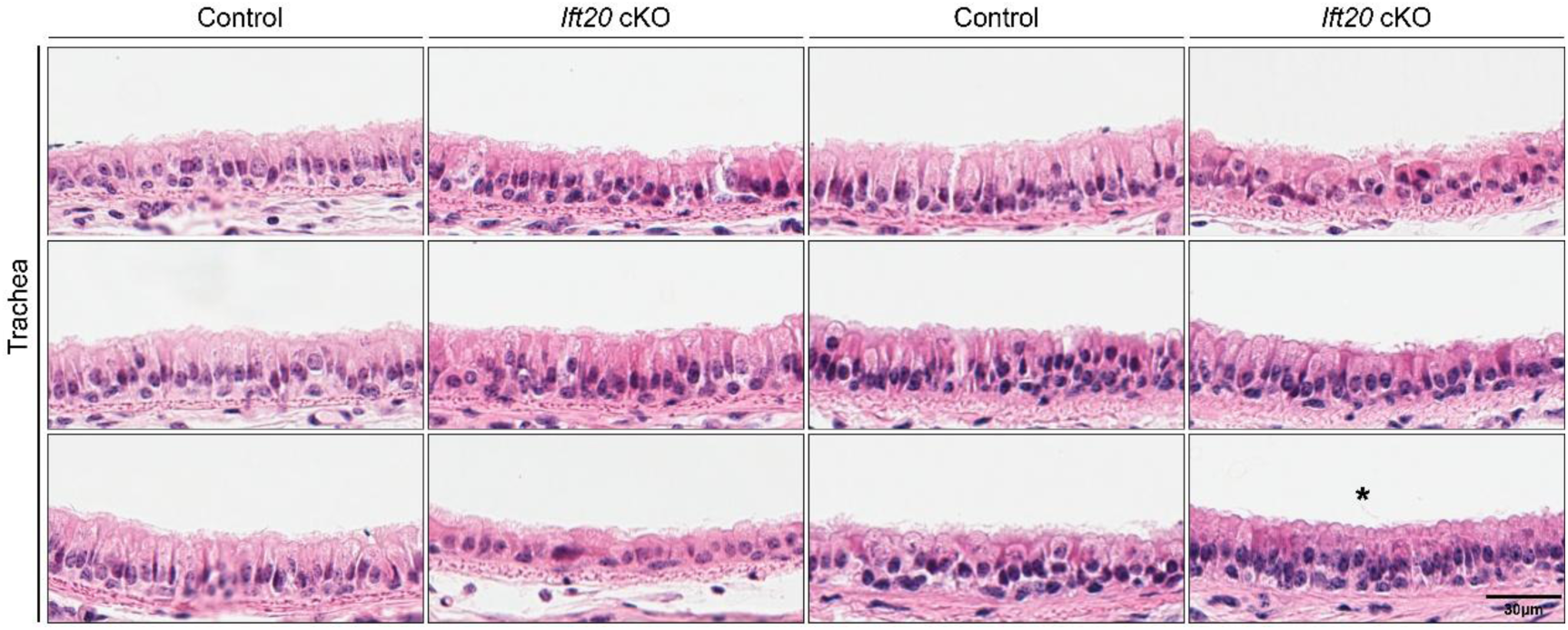
HE staining of tracheal sections from six pairs of mice. Motile cilia in some cKO mice showed no significant shortening. **\*** The image marked with an asterisk serves as a representative example of cKO mice where the motile cilia are noticeably shortened.

**Supplementary Figure 10.**
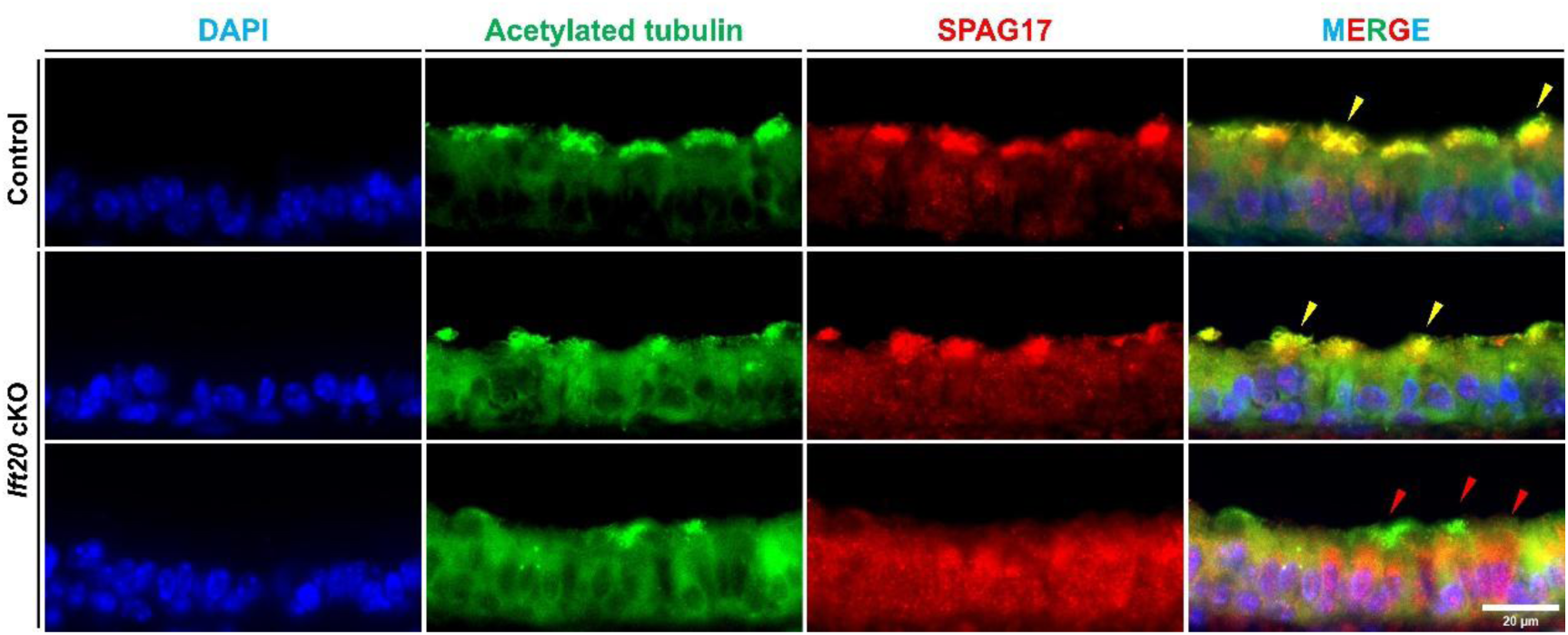
Immunofluorescence results of SPAG17 at 4 months of age. In the trachea of a cKO mouse, some SPAG17 was not transported to the motile cilia (red arrows), whereas yellow arrows indicate SPAG17 that was transported to the motile cilia.

**Supplementary Figure 11.**
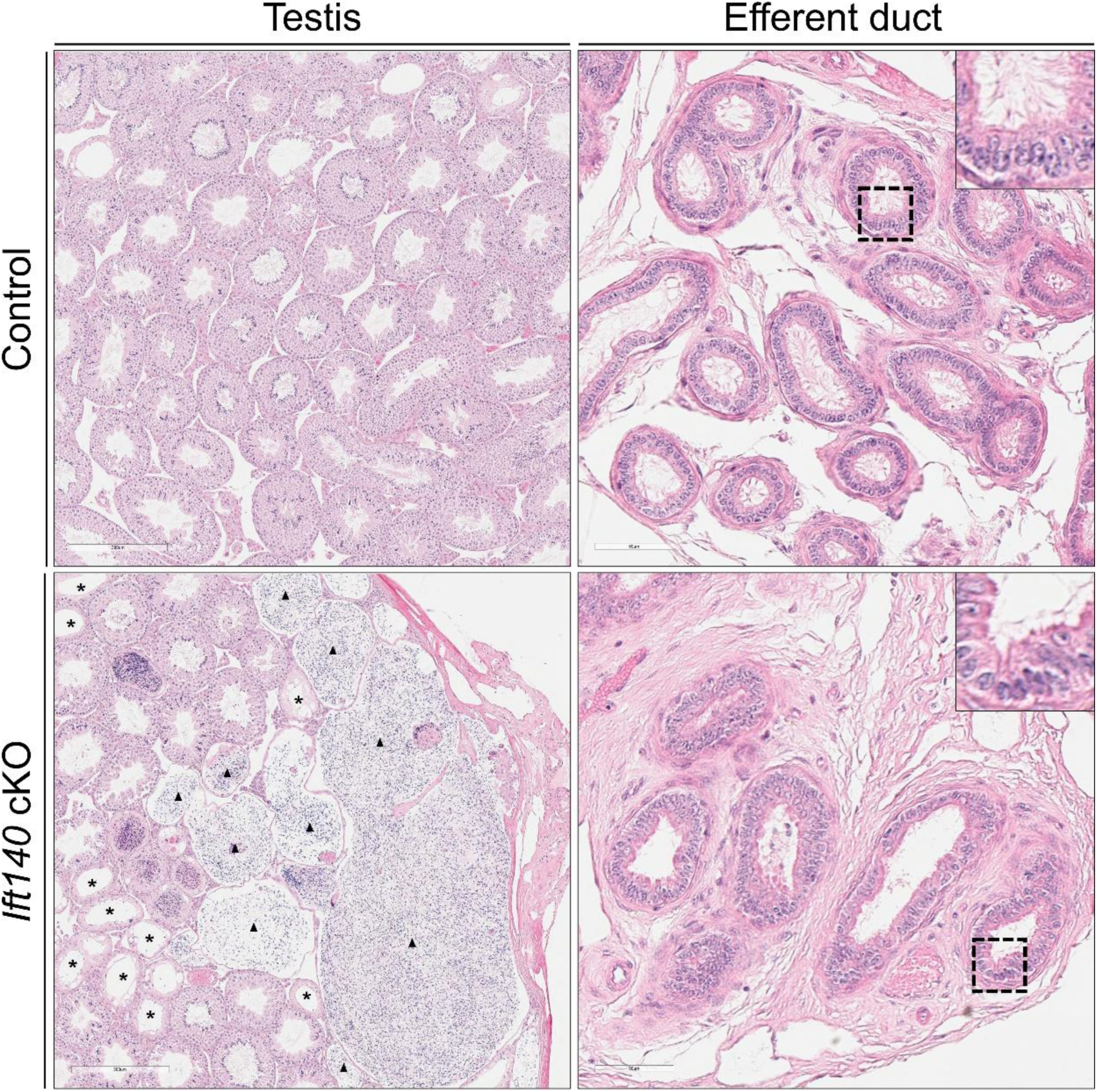
HE staining of the testes and efferent ducts in *Ift140-FOXJ1* cKO mice. Compared to the control group, the testes of cKO mice were significantly enlarged, with a large number of sperm obstructed in the rete testis and seminiferous tubules (marked by black triangles). Additionally, atrophy of the seminiferous epithelium in adjacent tubules was observed (marked by *). In the efferent ducts, the motile cilia of cKO mice were notably shortened.

**Table S1.** Antibodies used in this study.

| **Antibodies** | **Species** | **application** | **Concentration** | **Company source** | **Cat #** |
|---|---|---|---|---|---|
| IFT20 | Rabbit | IF | 1:200 | gift from Dr. Pazour |  |
| Acetylated-Tubulin | Mouse | IF | 1:200 | Proteintech | 66200-1-Ig |
| SPAG16 | Rabbit | IF | 1:300 | generated by our laboratory |  |
| SPAG17 | Rabbit | IF | 1:200 | generated by our laboratory |  |
| DNAI2 | Rabbit | IF | 1:300 | Proteintech | 17533-1-AP |
| DNAH2 | Rabbit | IF | 1:1000 | Sigma Aldrich | HPA067103 |
| Anti-Mouse IgG H&L -Alexa Fluor® 555 | Goat | IF | 1:4000 | Abcam | ab150114 |
| Anti-Rabbit IgG H&L-Alexa Fluor® 488 | Goat | IF | 1:4000 | Abcam | ab150077 |
| DAPI |  |  |  | Vector Laboratories | H-1200-10 |
| IFT20 | Rabbit | WB | 1:1000 | gift from Dr. Pazour |  |
| GAPDH | Mouse | WB | 1:1000 | Santa Cruz | sc-47724 |
| β-actin | Mouse | WB | 1:20000 | Proteintech | 66009-1-Ig |
| HRP-conjugated Goat Anti-Rabbit IgG(H+L) | Goat | WB | 1:2000 | Proteintech | SA00001-2 |
| HRP-conjugated Goat Anti-Mouse IgG(H+L) | Goat | WB | 1:2000 | Proteintech | SA00001-1 |

